# Guided assembly of multispecies positive biofilms targeting undesirable bacteria

**DOI:** 10.1101/2024.10.16.618781

**Authors:** Virgile Guéneau, Laurent Guillier, Cécile Berdous, Marie-Françoise Noirot-Gros, Guillermo Jiménez, Julia Plateau-Gonthier, Pascale Serror, Mathieu Castex, Romain Briandet

## Abstract

The use of synthetic microbial communities (SynComs) engineered to form positive biofilms that prevent the settlement of harmful bacteria is emerging as a promising strategy in biotechnology, particularly in reducing reliance on chemical antimicrobials. Despite this potential, the rationale for selecting specific strains in SynComs and the mechanisms underlying their antagonistic effects remains insufficiently understood. In this study, we present a bottom-up approach integrating live-cell imaging with high-throughput analysis of multi-strain biofilms across diverse scenarios. Through this method, we identified beneficial strains based on their superior ability to exclude undesirable bacteria and form mixed biofilms. Notably, our findings revealed that competitive strains against undesirable bacteria could also exclude other beneficial strains, emphasising the need for compatibility control in SynComs design. SynComs composed of *B. velezensis* and *Pediococcus* spp. demonstrated enhanced pathogen exclusion compared to single strains. Temporal analysis of biofilm interactions, supported by mathematical models, showed that pathogen exclusion was primarily driven by nutritional competition (Jameson effect) with additional specific interference mechanisms (prey-predator Lotka-Volterra model). Furthermore, pre-establishing SynComs to surfaces significantly increased pathogen inhibition, indicating a distinct biofilm-associated exclusion effect. These insights offer a framework for rational SynCom design and deepen our understanding of the mechanisms underpinning positive biofilm applications.

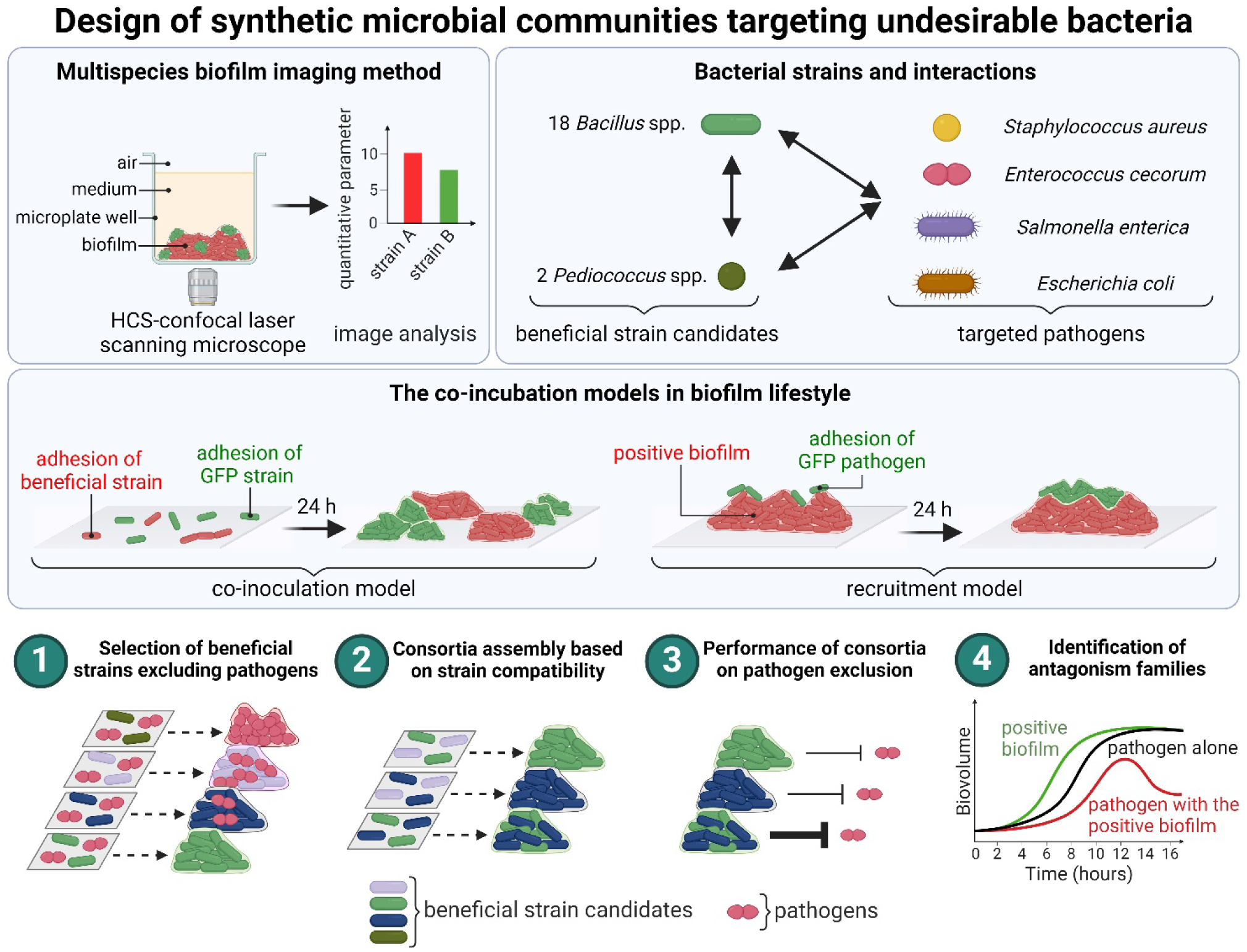

## Introduction

Microorganisms predominantly exist as biofilms, which colonise a wide variety of biotopes on Earth [1]. Biofilms are intricate communities of spatially organised microorganisms embedded in a self-produced matrix. These communities thrive at interfaces, interacting with each other and their environment [2]. The three-dimensional structures, coupled with diverse cell populations and matrix, endow biofilms with unique properties distinct from planktonic lifestyles [3–5]. Biofilms typically comprise multiple species engaged in various interactions, including resource competition, cooperation, and inhibition of other species [6,7]. Deciphering these complex interaction networks to predict biofilm behaviour and functions is a pivotal challenge in microbial ecology.

In line with the “One Health” concept to rationalise the use of chemical antimicrobials, biotechnological solutions are being developed to regulate interactions within complex biofilm communities through guided microbial ecology [8]. For instance, single strains or synthetic microbial communities (SynComs) [9,10] capable of forming positive biofilms with antagonistic activity against undesirable microorganisms can be intentionally introduced in the foods chain [11], directly onto hosts [12,13] but also more recently on surfaces such as livestock buildings [14].

A competitive ecological process can be established between bacterial species, resulting in the dominance of one and the exclusion of the other [15]. Pathogen exclusion can be attributed to nutritional [16] and spatial competition [17], as well as the secretion of interfering molecules through quorum sensing modulation [18], bacteriostatic [19], or bactericidal effects [20]. Each exclusion mechanism can be represented using mathematical modelling tools based on temporal analyses [21]. For instance, the Jameson-effect model can be described as competition between species over the use of environmental resources in order to maximise their growth and population. This model accounts for a nutritional competition which results in the deceleration of the population growth when the common resource(s) are exhausted [16]. Another model, Lotka-Volterra prey-predator model, implicates the secretion of interfering molecules, leading to the decline of the prey population. [22]. These models provide quantitative parameters describing the evolution of the partners and their mutual influence. However, quantifying antagonism typically relies on assessing planktonic interactions, which do not accurately reflect real conditions under which microbial communities reside in biofilms [23]. Furthermore, the inclusion of multiple strains in SynComs lacks clear justification and assurance of compatibility. In addition, the enhanced effects of SynComs compared to their constituent strains are often unclear.

To enhance the design of SynComs for positive biofilm applications, we developed a systematic pipeline to select beneficial strain candidates that specifically antagonise pathogens through their biofilm formation. SynComs were designed based on the ability of strains to coexist in the same positive biofilms without excluding partners. The objective was to identify compatible beneficial strains able to form a mixed biofilm with enhanced pathogen exclusion and biofilm formation capability compared to individual strains. This bottom-up approach relies on non-destructive observation of multispecies biofilm phenotypes using high-content screening confocal laser scanning microscopy (HCS-CLSM) combined with genetically engineered fluorescent strains and dedicated image analysis [24]. The study includes over 23 000 meticulously analysed z-stack images, predominantly utilising two colour channels to extract quantitative data on positive biofilm efficiency. The collection of tested beneficial strain candidates comprised 18 *Bacillus* and 2 *Pediococcus* strains. These genera are renowned for their biofilm-forming abilities [25,26], exclusion capabilities [20,27], and versatile applications in biotechnology, biocontrol [28] and biopreservation [11], making them attractive candidates for investigating their potential to create positive biofilms. The 20 candidate strains were screened for their impact on the growth and establishment of several pathogenic bacteria affecting people and/or animals, including *Staphylococcus aureus*, *Enterococcus cecorum*, *Escherichia coli*, and *Salmonella enterica* serovar Enteritidis (*S. enterica*), in two submerged mixed-species biofilm models developed for this study.

Through 4D (xyzt) HCS-CLSM imaging of interspecies interactions in biofilms, the underlying family of mechanisms driving antagonistic interactions on pathogens by the SynComs were elucidated. Moreover, we showed by modelling of biofilm interaction curves that these mechanisms depend on the initial quantities of each partner in the mixed biofilm and are specific to the biofilm lifestyle.

Together, these results allow for the rationalisation of SynCom formulation for positive biofilm application to limit pathogen growth and establishment on surfaces, while understanding the families of interactions involved.

## Materials and methods

### Bacterial strains and genetic constructs

The wild-type (WT) bacterial strains and genetic constructions used in the study are listed in **Sup. 1**. The phylogenetic analysis of the strains was conducted using the bacterial phylogenetic tree service provided by BV-BRC [29]. The phylogenetic tree was then constructed using the default codon tree method. For species assignments, the genome sequences were compared to those of the closest species within the *Bacillus* or *Pediococcus* genus in the BV-BRC database, using 1,000 genes with a tolerance for five deletions. Subsequently, the beneficial strains were ordered based on their phylogenetic distances (**Sup. 2**). The transformation protocol using the pCM11 plasmid derivatives, carrying the GFP or the mCherry-encoding genes, was adapted for each strain and is detailed in **Sup. 3**. Briefly, wild type *E. coli* and *S. enterica* were transformed using a standard heat shock protocol [30]. The protocols for preparing electrocompetent cells and transforming *E. cecorum* were adapted from Dunny *et al.* [31]. *B. velezensis* was transformed based on its natural competence [32] using a methodology inspired by Dergham *et al.* [33]. The plasmid stability assay is presented in **Sup. 4**.

### Biofilm models

All experiments were conducted at 30°C using Tryptic Soy Broth (TSB) medium (BioMérieux, France) supplemented with antibiotics (5 µg/mL erythromycin for Gram-positive species or 100 µg/mL ampicillin for Gram-negative species) when appropriate. 5 mL overnight cultures, inoculated from a glycerol stock at -80°C, were centrifuged at 5000 g for 5 minutes and then re-suspended in fresh TSB medium prior to conducting each experiment. The biofilms were cultivated at the bottom of a μclear® 96-well plate (Greiner Bio-one, France) which is compatible with high-resolution fluorescence microscopy. In this study, we have developed two biofilm models to study microbial interactions in multi-strain biofilms.

### Co-inoculation model

The co-inoculation model was developed with the objective of facilitating the controlled ratios of adhered strains on samples while limiting the presence of swimming planktonic cells. Multi-strain biofilms were established using fluorescent genetically labelled pathogenic strains in conjunction with non-labelled strains (one, two, or three strains). The initial adhesion biovolume between the fluorescent strain and the non-labelled strains was standardised using the protocol described by Guéneau *et al*. [24], which is based on a dual labelling by the GFP and the SYTO61, a cell-permeant red dye that binds to nucleic acid (Invitrogen, Carlsbad, CA, USA.). In this system, cells expressing the GFP will exhibit both green and red fluorescence, whereas untagged cells will only display the red fluorescence. In our kinetic measurements, the SYTO61 has been replaced by the FM4-64, a lipophilic red dye that binds to cell membranes (Invitrogen, Carlsbad, CA, USA) at a vital concentration of 1 µg/mL [34]. Three initial adhesion ratios were subjected to kinetic experiments. The first comprised two times more pathogens compared to the beneficial strain(s) (ratio 1). The second had two times more beneficial strain(s) compared to the pathogens. The third had 10 times the amount of the beneficial bacteria candidate(s) (ratio 3).

In practice, overnight cultures of the GFP-labelled and genetically unlabelled strains were diluted in 1 mL of TSB to achieve the desired adhesion ratio. 200 µL of the bacterial solution were added to the µclear 96-well plates and allowed to adhere statically at 30°C for 1.5 hours. The supernatant was then replaced with fresh TSB and the cultures were incubated for 24 hours at 30°C. The same protocol was employed to assess the compatibility between the GFP-labelled *B. velezensis* strains and the other beneficial strain candidates. The adhesion ratios were determined with the GFP-labelled *B. velezensis* 12048 and susbsequently applied to *B. velezensis* 11285 and *B. velezensis* ILPB8. In the kin consortium compatibility assay composed of these three *B. velezensis* strains, the SYTO9, a cell-permeant green dye that labels nucleic acid, was employed at a final concentration of 2 µg/m. In conjunction with the SYTO9, another nucleic acid labelling dye, DAPI was used (Invitrogen, Carlsbad, CA, USA). In this experiment, a LSM 700 inverted Confocal Scanning Laser Microscope (Carl Zeiss, Germany) equipped with a 405 nm laser was utilised to facilitate the excitation of the blue nucleic acid dye DAPI.

### Recruitment model

The recruitment model was developed for the purpose of examining pathogen adhesion and growth on a pre-established positive biofilm. The overnight cultures of pathogens were subjected to centrifugation at 5,000 rpm for 10 minutes, and re-suspended in fresh TSB medium in order to remove any residual antibiotics. In the case of the kinetics experiment, the resulting suspension was then supplemented with 1 µg/ml of FM4-64 membrane dye. A 50-microliter aliquot of a GFP-labelled pathogen suspension was added to wells containing a 24-hour-old positive biofilm and allowed to adhere statically at 30°C for 1.5 hours. Subsequently, the supernatants containing non-adhered cells were replaced with 200 µL of fresh TSB. CLSM acquisitions were conducted either directly (recruitment t=0h) or after 24 hours of growth at 30°C (recruitment t=24h). Prior to acquiring images via CLSM, 50 µL of a TSB solution containing SYTO61 at 2 µg/mL or FM4-64 at 1 µg/mL (for the kinetic assays) was added to the wells.

### Live-cell fluorescent imaging using CLSM

Live-cell fluorescent imaging was conducted using a Leica SP8 AOBS inverted high-content screening confocal laser scanning microscope (HCS-CLSM, LEICA Microsystems, Germany, MIMA2 platform of INRAE, https://doi.org/10.15454/1.5572348210007727E12). Biofilm images were acquired using a 63x water immersion objective with a numerical aperture of 1.2. Imaging was performed at a frequency of 600 Hz, with images taken every micrometre to ensure comprehensive coverage of the full height of the biofilm. The images had a resolution of 512 × 512 pixels, covering a physical area of 184.52 µm x 184.52 µm with a pixel size of 0.361 µm. For fluorescence detection, SYTO9 and GFP were excited with an argon laser at 488 nm, and the emitted fluorescence was collected using a hybrid detector (HyD LEICA Microsystems, Germany) in the range 500-550 nm.

FM4-64 was excited through the use of a helium-neon laser at 561 nm, and the emitted fluorescence subsequently collected with a hybrid detector in the range of 600-750 nm. Excitation of SYTO 61 was achieved at 633 nm and the emitted fluorescence was collected with a hybrid detector in the range of 650-750 nm.

For 4D (xyzt) acquisitions, the temperature was maintained at 30°C and 150 µm stacks were automatically acquired every hour using the HCS mode of the confocal microscope. Each experiment included between 3 and 6 biological replicates, with 4 technical replicates per biological replicate, thereby providing at least 12 technical values per condition.

### CLSM image analysis

Two-dimensional projections of biofilms and movies were generated using IMARIS 9.3.1 software (Bitplane, Zurich, Switzerland). Quantitative analysis of image stacks were performed using the BiofilmQ software [24,35]. Each fluorescence channel was analysed separately using the OTSU thresholding method and “global biofilm properties” were selected to extract the biovolume of the binarized images.

To assess the activity of *Bacillus* species strains against pathogens, the GFP biovolume (µm^3^/µm^2^) of submerged mixed biofilms was quantified and normalised to the biovolume of biofilms of a specific pathogen labelled with GFP. To standardise a score reporting biofilm anti-pathogenic activity, values were centre-reduced to fall between 0 (indicating the lowest activity) and 1 (representing the highest activity). This scoring method facilitated comparison of the anti-pathogenic activity of different *Bacillus* strains, allowing the identification of the most promising potential.

### Modelling microbial growth and competition

GFP-measured biovolumes served as model inputs for both pathogens. For *Bacillus* in monoculture, biovolumes corresponding to the FM4-464 marker were used. In co-culture, beneficial strain(s) biovolumes were determined by subtracting the GFP-biovolume of the co-inoculated pathogen, from the total biovolume.

For monoculture experiments, biovolume increase was described using a generic primary growth model [36] :

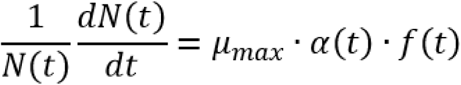

This model defines *µ*_max_ as the exponential growth rate, α(t) as the adjustment function, and *f*(t) as the inhibition function. The nlsMicrobio package was employed to fit the Baranyi & Roberts primary growth model, providing estimates for *µ*_max_, lag phase (derived from α(t)), maximum population size (*N*_max_, derived from f(t)), and initial population size (*N*_0_).

Two types of models were fitted to the competition data between pathogens and bioprotective flora.

The first type of model is the Jameson-type model [37]. The Jameson effect can be described as competition between species to use environmental resources in order to maximise their growth and population. When the common resource(s) are exhausted, the competition is over and the growth of each species in the population stops.

The growth stops simultaneously by both populations. Both competitors share the inhibition function (*f*(t)) of the exponential growth.

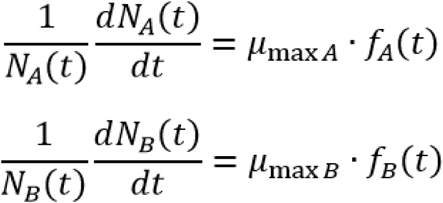

The inhibition function being

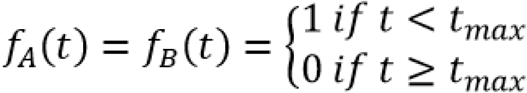

where *t*_max_ is the time at which the stationary phase begins for populations A and B.

The second type of model is the Lotka-Volterra model. In this model, the inhibition functions can be described as follow:

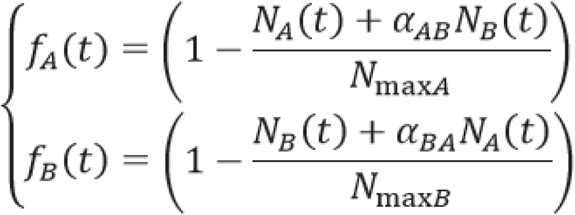

where the parameters *α_AB_* and *α_BA_* are the coefficients of interaction measuring the effects of one species on the other.

It makes no specific assumptions about the mechanisms underlying species interactions; they can be parameterised in ways that approximate any combination of underlying mechanisms. For example, in a system with two species, if both *α*_AB_ and *α*_BA_ are less than zero, species suppress each other’s growth, indicating competition.

### Statistical analysis

Results related to the description of biofilm phenotypes were presented as mean and standard deviation (SD). Statistical analysis was performed using two-way ANOVA followed by Fisher’s least significant difference without correction, utilising PRISM software (GraphPad, USA, California). Statistical significance was determined at a *p*-value less than 0.05. The significance levels are denoted as follows: * for *P <* 0.05, ** for *P* < 0.01, *** for *P* < 0.001, **** for *P* < 0.0001. The nlsMicrobio [38] and gauseR [39] packages were respectively used to fit the Barnayi&Roberts growth model, Jameson-type and Lotka-Volterra models to biovolumes.

## Results

### Selection of beneficial strains for pathogen exclusion

A collection of 20 beneficial strains from various isolation origins was first characterised for different surface-biofilm and motility phenotypes. These phenotypes included the formation of macro- colonies, the development of submerged biofilm, and the ability of strains to swarm at the surface of semi-solid agar. This phenotypic characterization revealed a wide range of phenotypic diversity (**Sup. 5**). While differentially affected by the ability to spread on solid (macrocolony) or semi solid (swarming) surfaces, all strains were observed to develop submerged biofilms of varying thickness and structures. This observation led us to use the submerged biofilm as a working model in our study. A screening process was conducted to identify beneficial strains capable of restricting the growth and establishment of four pathogens, *E. cecorum*, *S. aureus*, *E. coli* and *S. enteritica* (**Fig. 1A, B**). To this end, we developed two co-incubation biofilm assays. These assays were based on the formation of mixed biofilms after the co-inoculation of strains at different ratios and on the potential recruitment of a pathogenic strain by a preformed biofilm composed of potentially beneficial bacilli strain(s). To ensure a balanced initial co-inoculation of both the beneficial and pathogen strains, the adhesion ratio was first validated by CLSM (**Sup. 6**). After co-inoculation and 24 hours of growth, the raw biovolume of GFP-labelled pathogens from z-stacks were extracted by image analysis (**Sup. 7**). Regarding the recruitment assay, the raw biovolumes of the pathogen were quantified to investigate its adhesion (recruitment t=0 h, **Sup. 8**) and growth (recruitment t=24 hours, **Sup. 9**) on an established positive biofilm. The data were used to calculate antagonistic scores for each beneficial strain against pathogens (**Fig. 1C**). Our results revealed different modes of interactions between the beneficial bacilli and the pathogen. For instance, in a co-inoculation assay, *E. coli* and *S. enterica* were not excluded. Conversely, in the recruitment assay, the *B. velezensis* strains as well as *Paenibacillus* spp. 1167 exhibited a strong capacity to exclude *S. enterica*. Pathogen adhesion to pre-established positive biofilms (*i.e.*, recruitment at T=0 h) was reduced with most beneficial strains, except for *S. enterica*. Overall, *B. velezensis* consistently demonstrated superior pathogen exclusion capacity across the two interaction models in comparison to other beneficial strains. Remarkably, *B. velezensis* strains 11285, 12048, and ILPB8 consistently achieved the highest scores and illustrated superior performance against *E. cecorum* and *S. aureus*.

**Figure 1:**
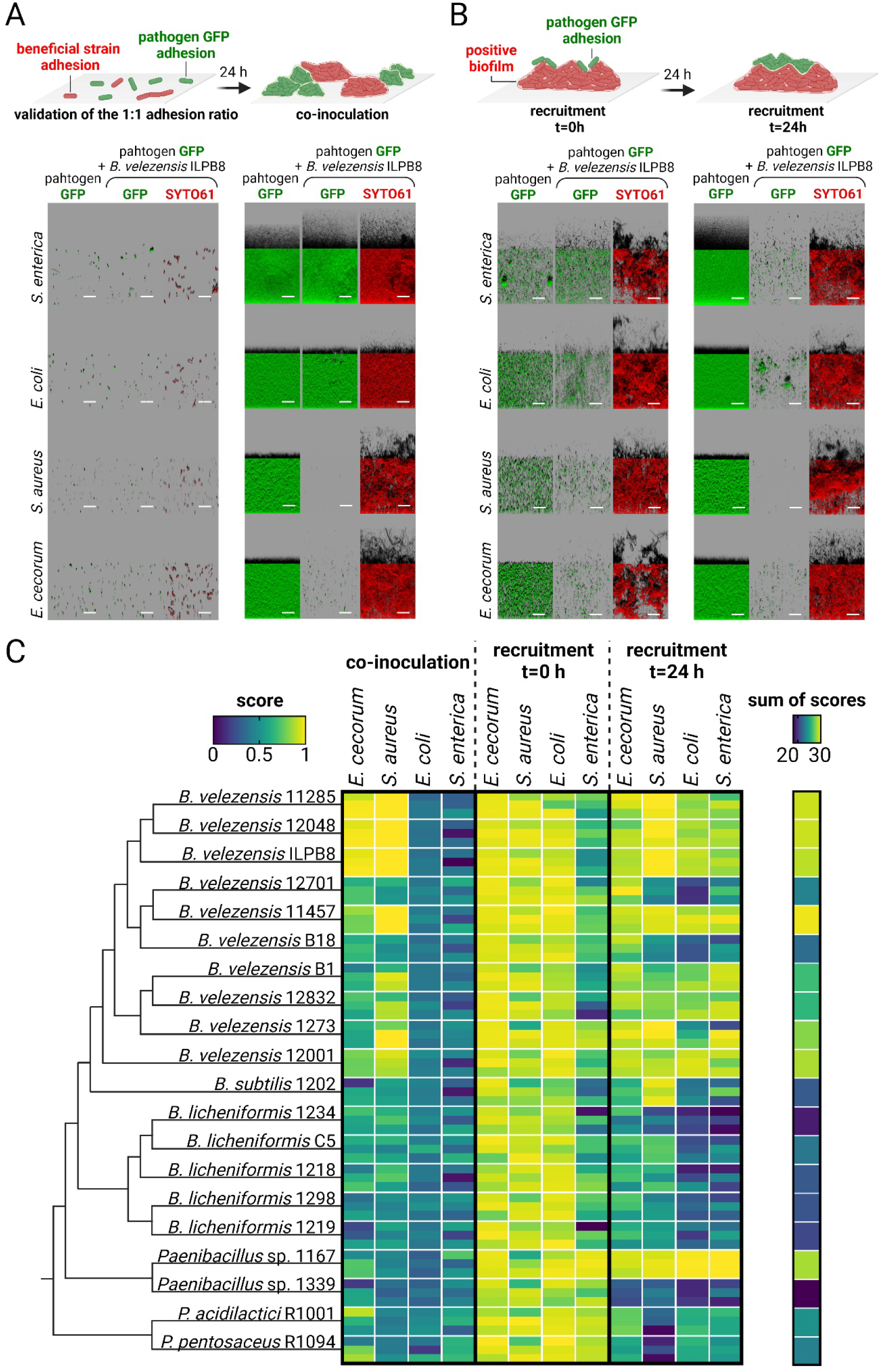
Antagonism scores of the candidate beneficial strains against pathogens in two co-incubation models. (A) In the co-inoculation model, adhesion ratios (0.50 +/- 0.25) of both pathogenic and beneficial strains were determined by image analysis of CLSM observations. Following adhesion with these pre-determined ratios, biofilms were incubated 24 hours before being observed with CLSM. (B) In the second model, a planktonic culture of the pathogen was added to a pre-established biofilm for 24 hours prior CLSM observation. After adhesion, a 24-hour incubation period was followed by CLSM observation. In both models, SYTO61 was used to visualise the entire population in red (i.e. pathogens and beneficial strains). Pathogen GFP biovolumes were quantified to identify beneficial strains that reduced pathogen biovolume compared to pathogens growing alone under the same conditions. Representative images are shown from the strain *B. velezensis* ILPB8. Scale bar = 40 µm. (C) Pathogens were grouped by model and the 20 candidate beneficial strains were ordered according to a phylogenetic tree reflecting their relatedness. In the heat map, a score of 1 represents the highest antagonist activity and 0 represents the lowest, calculated based on pathogen GFP biovolume in the presence of a beneficial strain normalised to the GFP biovolume of the pathogen growing alone. Each square represents one interaction, displaying the means of three biological replicates, each calculated from four technical replicates. The sum of scores against each pathogen across all models was calculated to determine the total antagonistic power of each beneficial strain.

Cell counts were performed using *B. velezensis* ILPB8 strain, which exhibits a high antagonistic score, thereby confirming that the reduction in the GFP signal in our experiments was a direct result of a decline in cell numbers (**Sup. 10**).

### Engineering a mixed biofilm from compatible beneficial bacterial strains

We selected *B. velezensis* strains 11285, 12048, and ILPB8 based on the highest sum of antagonist scores against pathogens. We employed a co-inoculation model to identify compatible mixed biofilms resulting from interactions between the three genetically GFP-tagged *B. velezensis* strains and other beneficial candidate strains. Initially, the adhesion ratios between *B. velezensis* 12048 GFP and 20 beneficial strains were determined and applied subsequently to *B. velezensis* 11285 GFP and *B. velezensis* ILPB8 GFP (**Sup. 11**). After 24 hours of growth, the biovolumes were extracted from the z-stacks of images. The GFP/SYTO61 ratio was calculated to assess the strains’ ability to coexist or exclude each other (see **Fig. 2A**, **B**). Our results suggest that beneficial strains exhibit mutual exclusion tendencies. Mixed biofilm formation occurred only when the three GFP-labelled *B. velezensis* strains were combined with their genetically unlabelled counterparts or with the two phylogenetically more distant *Pediococcus* spp. Notably, none of the combinations exhibited a total biovolume (SYTO61) significantly different from that of either strain when grown individually (**Sup. 12**).

**Figure 2:**
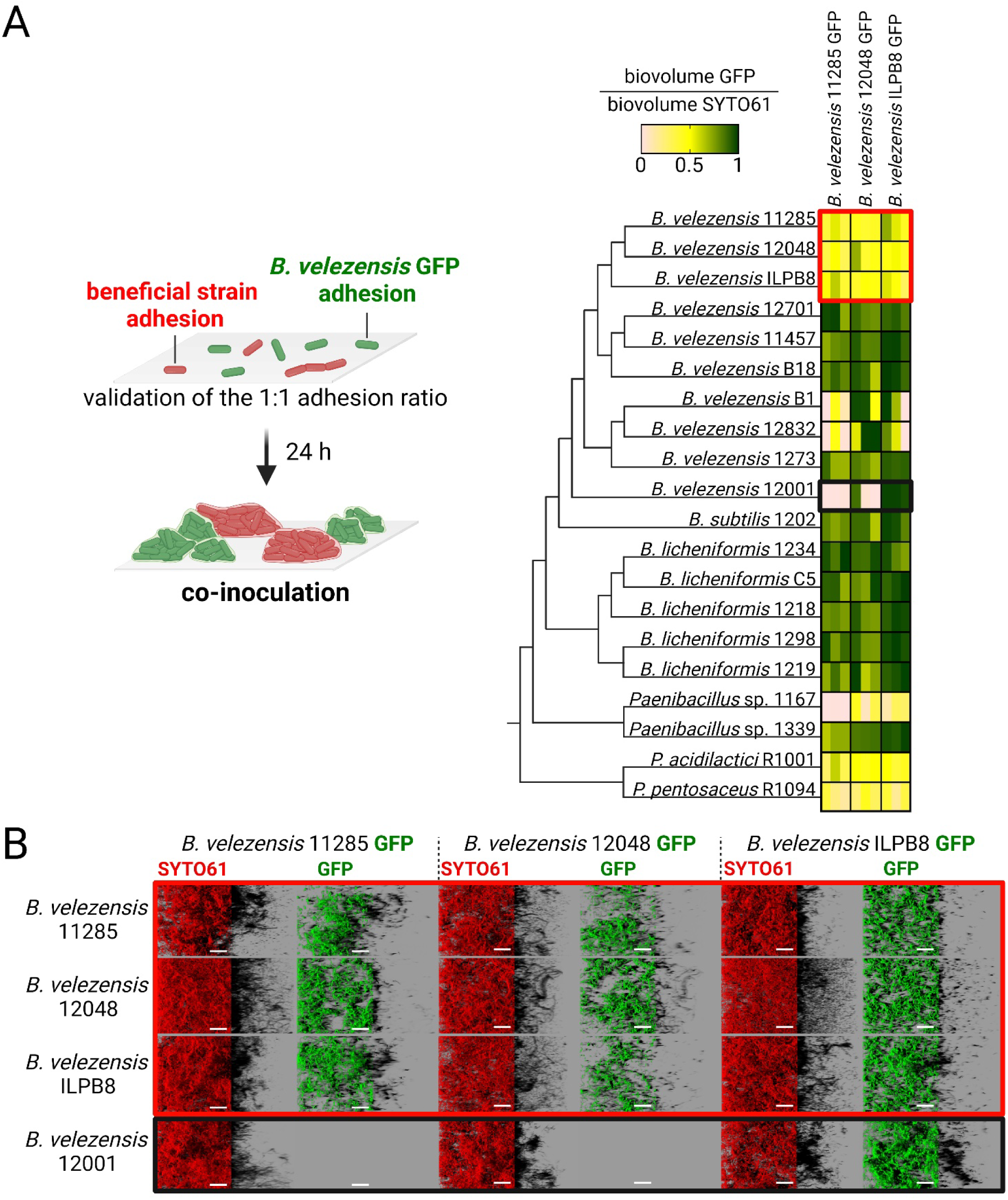
Mixed biofilm compatibility experiment between candidate beneficial bacterial strains. (A) The ability of three *B. velezensis* strains, expressing GFP and exhibiting high antagonistic activity against pathogens, to form mixed submerged biofilms with other beneficial strains was evaluated using the co-inoculation model. Strains were ordered according to their phylogenetic distance. The SYTO61 and GFP signals were quantified to calculate the GFP biovolume to SYTO61 biovolume. Each square represents one interaction and shows the means of three biological replicates, each derived from four technical replicates. Red and black rectangles highlight the examples illustrated below. (B) Representative examples illustrate the interactions between the three *B. velezensis* GFP strains capable of forming two-strain mixed biofilms. The exclusion scenario is demonstrated by *B. velezensis* 12001, which excludes *B. velezensis* 11285 and *B. velezensis* 12048, while it is excluded by *B. velezensis* ILPB8. Scale bar = 40 µm.

### A three-strain B. velezensis kin-compatible consortium covers more surface area but do not exhibit greater antagonism than single strains

Further studies were carried out to assess the potential of the selected *B. velezensis* strains 11285, 12048 and ILPB8, to form a three-strain mixed biofilm. Co-inoculation experiments were performed, incorporating two of the *B. velezensis* strains, genetically tagged with GFP or mCherry, together with the wild-type of the third *B. velezensis* strain (**Fig. 3A**). After staining the consortium with the DNA binding fluorescent dye DAPI, the observations and proportion quantifications of the three strains revealed a uniformly mixed biofilm (**Fig. 3B**). The strains within the kin consortium showed no significant differences in their biovolume (**Fig. 3C**). However, the kin consortium was found to exhibit a significantly higher surface area coverage than the one- and two-strains combinations (**Fig. 3D**).

**Figure 3:**
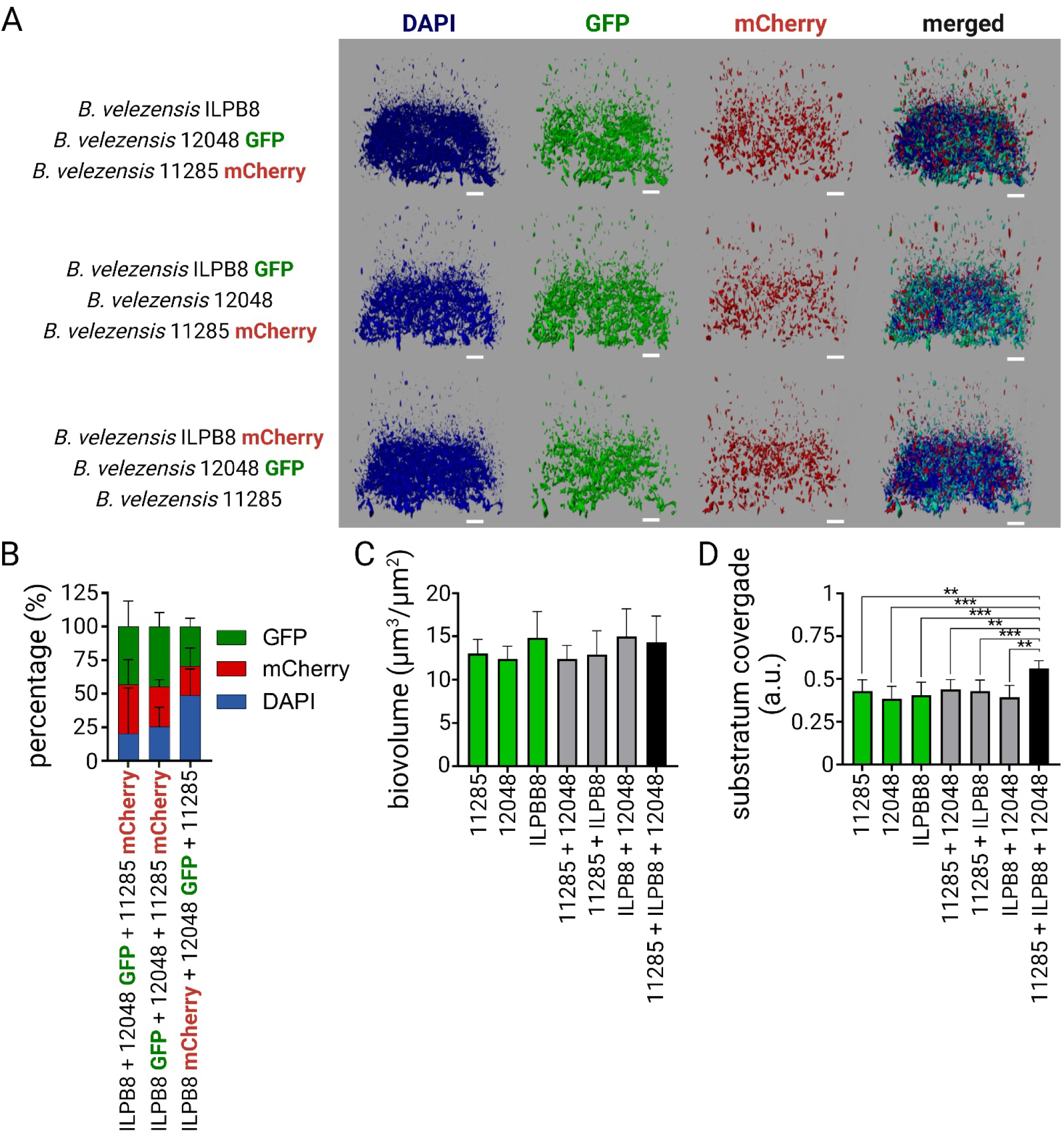
Mixed biofilm composed of three kin-compatible strains of *B. velezensis*. (A) Representative images of mixed biofilms using *B. velezensis* ILPB8 strains wild type, or genetically labelled with GFP or mCherry, along with the addition of DAPI in the biofilm to visualise the entire population in blue. Scale bar = 30 µm. (B) The percentage of signal from GFP-, mCherry- and DAPI-labelled cells relative to the total biovolume of DAPI in biofilms of the three strains is presented. (C) The co-inoculation model was employed using each wild-type strains, either alone or in combination, and the biofilms were labelled with SYTO9 and observed using the Leica SP8 HCS-CLSM to investigate their overall structure. Biovolume data were extracted and are shown in the graph. (D) Substratum coverage quantification of wild-type strains alone or in consortia is presented. Error bars correspond to standard deviation.

Further exploration of the mechanisms underlying the exclusion of *E. cecorum* and *S. enterica* by *B. velezensis* was conducted using HCS-CLSM kinetic experiments applied to the two co-incubation models. (**Sup. 13**). To achieve this, the vital membrane-dye FM4-64 was used to visualise the entire biofilm without altering the growth of the pathogen (**Sup. 14**). Biovolume curves over time were extracted from the images and processes using the Jameson or Lotka-Volterra mathematical models to determine the growth rate and the growth potential (**Sup. 15**). Additionally, to enhance our understanding of interaction mechanisms, distinct initial adhesion ratios of *B. velezensis* and pathogens were applied. The first ratio (ratio 1) comprised twice the number of pathogens in comparison to the beneficial strain(s). The second ratio (ratio 2) contains twice more beneficial strain(s) compared to the pathogens. Finally, the third ratio (ratio 3) has 10 times the amount of the beneficial bacteria candidate(s). We observed that the growth rate of *E. cecorum* was reduced in the presence of *B. velezensis* under all conditions except when the initial ratio started with a higher proportion of the pathogen (ratio 1) (**Fig. 4A**). For all ratios, the growth potential was found to decrease and even to become negative with the ratio 1, indicating a decline of *E. cecorum* biovolume at the end of the kinetics compared to the initial situation (**Fig. 4B**). In this specific case, the interactions were found to adhere to a Lotka-Volterra model. The growth rate and growth potential of *S. enterica* in the presence of *B. velezensis* were only observed to be altered in the context of recruitment (**Fig. 4C, D)**. With the exception of the ratio 1 in co-inoculation with *E. cecorum*, the interaction profiles were found to adhere to a Jameson model for all the strains of *B. velezensis* and their associated consortia against pathogens. However, significant differences in exclusion capacity between the *B. velezensis* strains against the two pathogens could be observed (**Sup. 16-17**).

**Figure 4:**
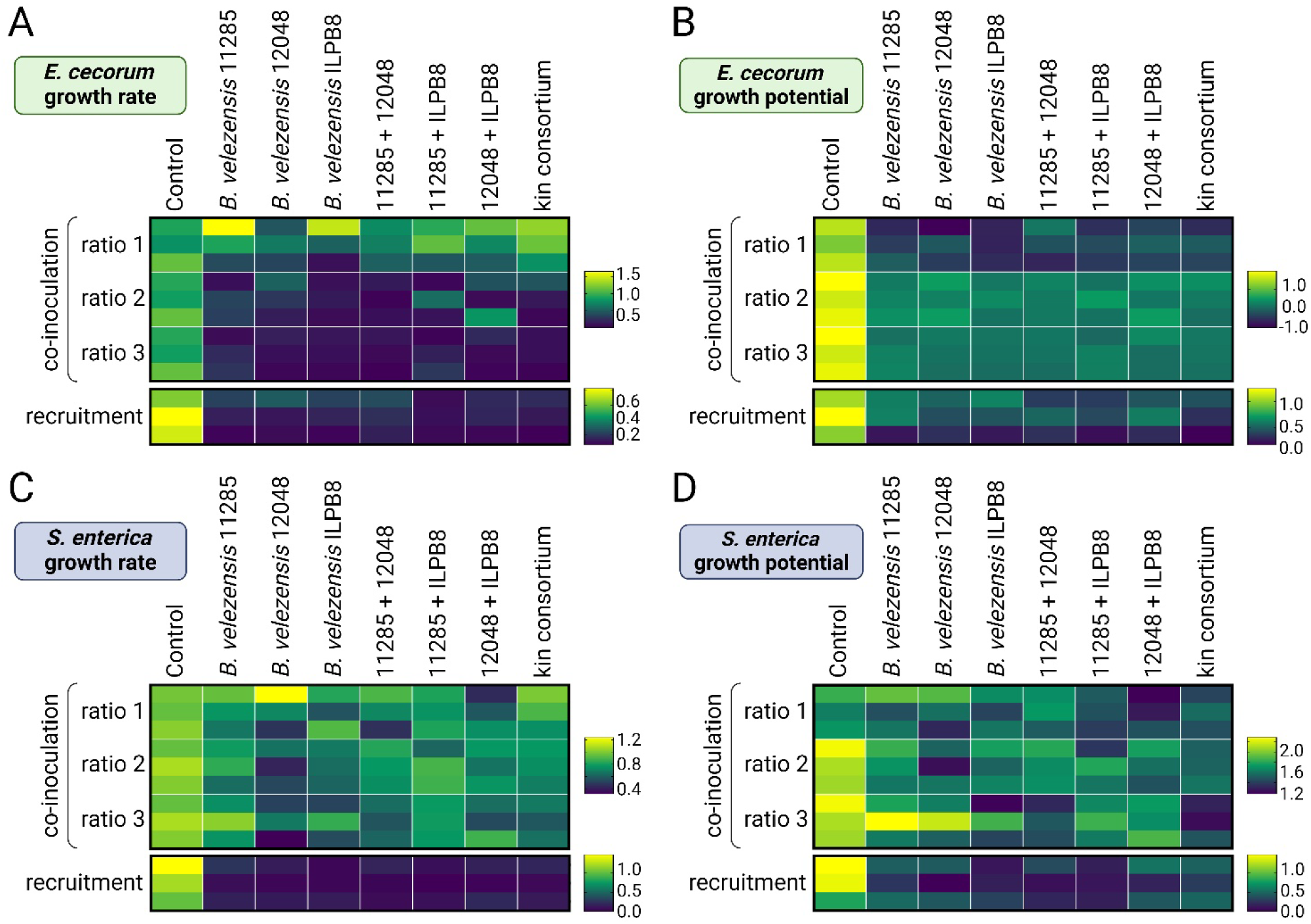
Pathogen growth parameters in mixed biofilms with *B. velezensis*. (A) Growth rate (h^-1^) determination of *E. cecorum* by modelling GFP biovolume curves and (B) calculation of growth potential for *E. cecorum* determined by subtracting the initial biovolume N0 from the final biovolume Nf (calculated in log10 (biovolume µm^3^/µm^2^)). The initial biovolume ratios of *E. cecorum* GFP to *B. velezensis* were determined at the start of the experiment (ratio 1 = 1.4 (+/- 0.2), ratio 2 = 0.3 (+/- 0.06), ratio 3 = 0.03 (+/- 0.02), recruitment = 0.2 (+/- 0.04)). (C) Growth rate determination of *S. enterica* by modelling GFP biovolume curves and (D) calculation of growth potential for *S. enterica* determined by subtracting the initial biovolume N0 from the final biovolume Nf (calculated in log10 (biovolume µm3/µm2)). The initial biovolume ratios of *S. enterica* GFP to *B. velezensis* were determined at the start of the experiment (ratio 1 = 3.2 (+/- 0.8), ratio 2 = 0.4 (+/- 0.1), ratio 3 = 0.1 (+/- 0.05), recruitment = 4.8 (+/- 0.8)). The kin consortium corresponds to the SynCom of *B. velezensis* strains 11285, 12048 and ILPB8. Each square represents one interaction performed using the HCS mode of the CLSM and shows the means of three biological replicates, each calculated from four technical replicates.

### Additive anti-pathogenic effects of B. velezensis and Pediococcus spp. compatible consortia

We demonstrated that the Kin consortium could form compatible and homogeneous mixed biofilms with the two *Pediococcus* strains *P. acidilactici* R1001 or *P. pentosaceus* R1094 **(Fig. 5 A, B)**. The four-strain consortia covered over 80% of the surface, mainly due to the influence of *Pediococcus* spp. biofilm, which developed between the clusters formed by the Kin consortium **(Fig. 5 A, C).** This biofilm exhibited a greater biovolume compared to controls **(Fig. 5 D)**.

**Figure 5:**
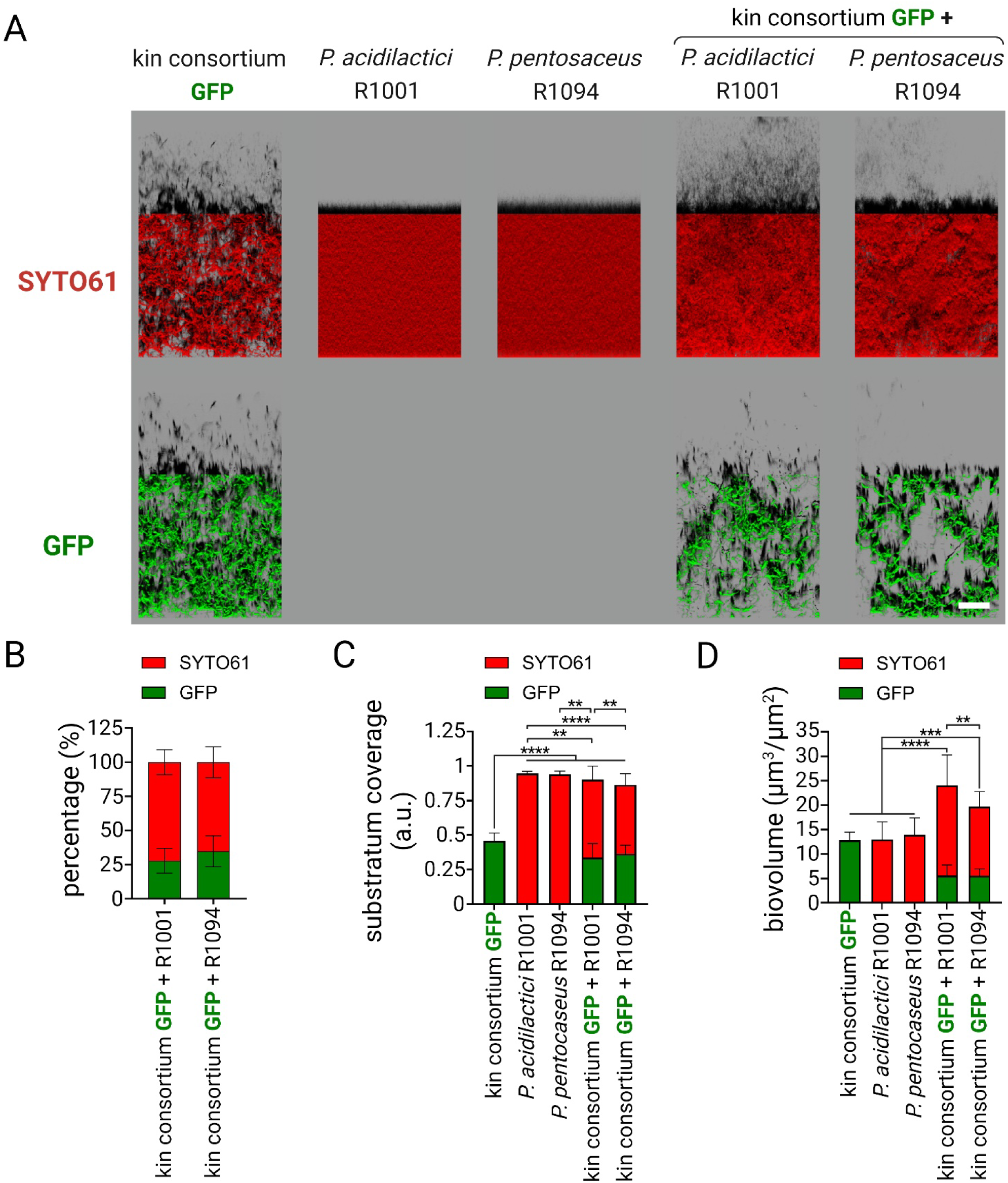
Compatible mixed biofilm composed of the kin consortium of *B. velezensis* with *Pediococcus* spp. (A) Representative images showing three kin-compatible strains of *B. velezensis* GFP (kin consortium: strains 11285 GFP, 12048 GFP and ILPB8 GFP) co-cultured together with *Pediococcus* spp. SYTO61 was used to label all the population in red, while *B. velezensis* were marked with GFP. Scale bar = 50 µm. (B) Percentage of signal from GFP and SYTO61, indicating the proportion of each strain within the biofilm (C) The co-inoculation model was used to investigate the overall structure of the biofilm. Biovolume data were extracted and are presented in the graph. (D) Substratum coverage quantification of strains alone or in consortia, illustrating the extent of surface coverage by each configuration. Error bars represent standard deviation.

The most effective exclusion activity against pathogens at 24h is achieved by mixed biofilms composed of *Pediococcus* spp. and *B. velezensis* strains from the Kin consortium (**Sup. 18 A**). Indeed, the consortia composed of *B. velezensis* and *Pediococcus* spp. demonstrated a consistently enhanced ability to exclude *S. enterica* at 24 hours compared to that of individual beneficial strains in the recruitment model (*P*<0.0001). In addition, the consortia performed at least as well as the best individual strains (**Sup. 18 B-G**). Therefore, we investigated the mechanisms of *S. enterica* exclusion in the first 12h of recruitment using HCS-CLSM coupled with modelling **(Fig. 6 A)**. The growth rate of *S. enterica* was found consistently reduced in the presence of beneficial strains, with no improvement observed in the Kin consortium when combined with *Pediococcus* spp. **(Fig. 6 B)**. However, a significant (*P*<0.01) decrease in *S. enterica* growth potential was observed when *P. pentosaceus* R1094 was added to the Kin consortium, compared to the performance of either the Kin consortium or *P. pentosaceus* R1094 alone **(Fig. 6 C)**.

**Figure 6:**
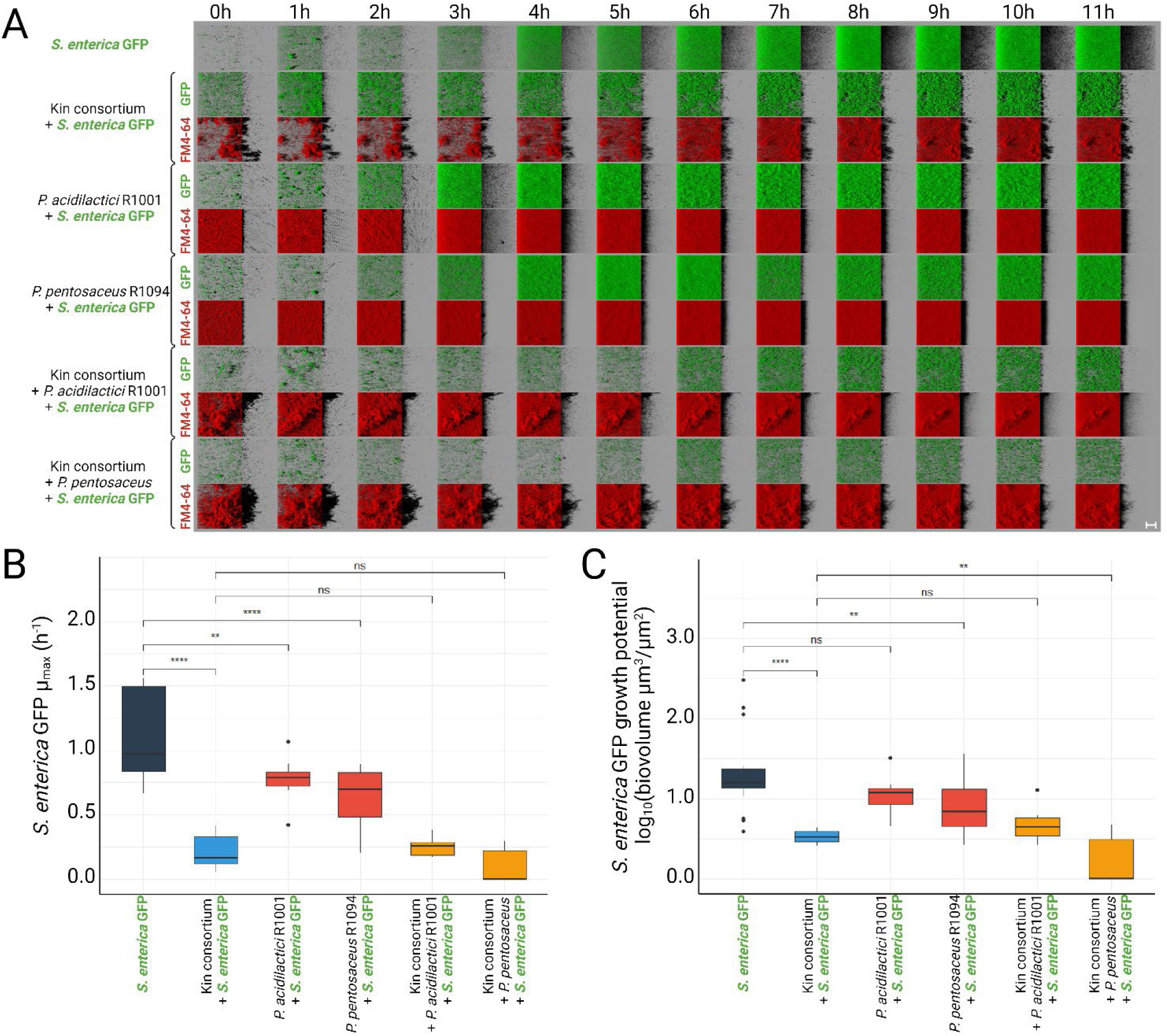
Characterisation of antagonistic activity against pathogens by consortia of *B. velezensis* and *Pediococcus* spp. in the recruitment model. (A) Representative CLSM images showing biofilm growth over time for each condition. The green colour corresponds to GFP labelling of *S. enterica*, while the red colour represents total population labelling by FM4-64. The SynCom of *B. velezensis* strains 11285, 12048 and ILPB8 (kin consortium) was co-cultured with *P. acidilactici* R1001 or with *P. pentosaceus* R1094 before the adhesion of *S. enterica* GFP in the recruitment model. The initial biovolume ratios of *S. enterica* GFP to the beneficial strains were determined at the start of the experiment (Kin consortium = 6.3 (+/- 2.2), R1001 = 0.14 (+/- 0.05), R1094 = 0.44 (+/- 0.20), Kin consortium + R1001 = 0.20 (+/- 0.03), Kin consortium + R1094 = 0.40 (+/- 0.09)). Scale bar = 40 µm. (B) Box plot modelling of *µ*_max_ of *S. enterica* GFP in the recruitment model, showing statistical analysis of the additive effects of *B. velezensis* and *Pediococcus* spp. consortia (x + y) on pathogen growth inhibition. Error bars represent standard deviation. (C) Box plot modelling of growth potential of *S. enterica* GFP in the recruitment model, showing statistical analysis of the additive effects of *B. velezensis* and *Pediococcus* spp. consortia (x + y) on pathogen growth inhibition. Error bars represent standard deviation.

We investigated whether the acidification of the environment by *Pediococcus* spp. was involved in enhancing the effect of *B. velezensis* on the exclusion of *S. enterica* in the recruitment t=24h model (**Sup. 19**). We demonstrated that acidification proportionally reduced the biovolume of *S. enterica* biofilms but did not affect the exclusion by *B. velezensis* ILPB8 (**Sup. 20**).

Interestingly, the biofilm lifestyle influences interactions differently compared to the planktonic lifestyle. Indeed, in the planktonic mode, *Pediococcus* spp. are excluded by *B. velezensis* (**Sup. 21**). *S. enterica* systematically excludes *B. velezensis* or *Pediococcus* spp. when starting with the same ratio of the two partners in planktonic cultures. Furthermore, starting with the same initial inoculum as for recruitment, but agitating the cultures under planktonic conditions, a culture containing *B. velezensis* is able to completely inhibit the growth of *S. enterica*, whereas *Pediococcus* spp. alone was excluded (**Sup. 22**).

## Discussion

This study presents a rational and systematic approach for selecting and combining strains within SynComs to form stable biofilms with robust antagonistic activity against bacterial pathogens. By employing a non-destructive high-throughput imaging pipeline, we demonstrated that compatible strains, such as *B. velezensis* and *Pediococcus* spp., can form stable mixed biofilms with enhanced pathogen exclusion capabilities. These findings underscore the potential application of these communities in positive biofilm strategies for surface microbial management.

Common methods for studying microbial interactions in multi-species communities, such as metabarcoding [40], optical density measurements [41], or colony-forming unit (CFU) counting [42], often disrupt intricate three-dimensional structures crucial in biofilm studies [43]. In contrast, our approach utilised non-destructive imaging techniques, allowing precise quantification of individual community members and observation of their spatial arrangement over time. This approach addresses significant gaps in existing methodologies, which often neglect critical factors such as adhesion rates and the formation of filamentous structures or aggregates [2,25]. Our co-inoculation model, which used image-based post-adhesion biovolume calibration, ensured accurate initial quantification of both partners on the substrate - a crucial step in understanding biofilm dynamics, where spatial arrangements strongly influence microbial interactions [44].

Despite the advantages of microscopy techniques, including their capacity for high throughput and detail analysis, they are inherently time-consuming and generate large volumes of data. Furthermore, distinguishing the contributions of each strain within biofilms composed of more than three species presents significant challenges. To overcome these hurdles, genetic manipulation of strains for the expression of distinct fluorophores or the use of specific dyes is necessary. The selection of fluorophores with distinct absorption and emission spectra is critical to prevent overlap within the same sample. These technical limitations restricted our ability to explore all potential strain combinations, such as co-culturing *P. acidilactici* and *P. pentosaceus,* due to the absence of corresponding fluorescent strains. Future investigations should consider the microbiota context of the studied ecosystem, as it may influence observed interaction phenomena [45].

Through temporal imagining and mathematical modelling, we identified key mechanisms driving pathogen exclusion within biofilms, primarily involving spatial and nutritional competition, along with the potential synthesis of antimicrobial compounds. Our findings suggest that pathogen exclusion is largely governed by the Jameson effect, coupled with specific interference mechanisms, where both the growth potential (nutritional competition) and the growth rate (secretion of bacteriostatic molecules) of the pathogen are reduced [16]. While this image-based approach provides a general framework for understanding these dynamics, it does not pinpoint specific molecular effectors responsible for pathogen exclusion. Future studies using omics approaches in combination with genetic analysis may help to elucidate these molecular underpinnings of pathogen exclusion [46].

The inhibitory effect of *B. velezensis* on a broad range of bacteria is well documented, often attributed to the secretion of interfering molecules [47], some of which are specifically expressed in biofilms [48]. In interactions that align with the Lotka-Volterra prey-predator model, mutual influence between microbial partners is observed [22]. Interestingly, at certain concentrations, the positive biofilm would secrete a molecule capable of killing *E. cecorum,* although the specific nature of this molecule remains unknown. These findings emphasise the need to consider the biofilm lifestyle, as their related kinetics differ significantly from those in planktonic cultures, even when starting with the same initial ratios of beneficial strains and the evaluated pathogen. This distinction is crucial, as metabolic processes vary widely between biofilm and planktonic states [5]. Therefore, the biofilm lifestyle should be a key consideration in designing SynComs intended for environments where bacteria are likely to form biofilms. Moreover, the experiments were conducted under controlled laboratory conditions with a maximum of 5 partners. A future perspective of this work would be to consider the microbiota of the specific surfaces to be treated in order to study the interactions, which is important to take into account, as it can potentially exclude both pathogens and intentionally applied beneficial SynComs [49].

Before evaluating the antagonistic effects of SynComs on pathogen growth, we systematically assessed the compatibility of strains to establish stable mixed biofilms without excluding other partners. Our results revealed that *B. velezensis* strains, selected for their ability to exclude pathogens, also tend to exclude other *Bacillus* spp. However, the compatibility between the three *B. velezensis* strains is consistent with the principle of kin discrimination, whereby organisms differentiate between genetically related (kin) and unrelated (non-kin) individuals [50,51]. This discrimination is likely based on the detection of small signalling molecules or specific flagellin motifs in *B. velezensis* [52].

Notably, only the three competitive *B. velezensis* strains were able to form mixed biofilms with one another and with the more distantly related *Pediococcus* spp. Even so, no significant increase in pathogen exclusion was observed in these kin-compatible consortia compared to individual strains, likely due to the presence of shared competition mechanisms and substantial overlap in gene content. By selecting more phylogenetically distant but compatible species, such as *B. velezensis* and *Pediococcus* spp., we have achieved genetic diversification, enhanced exclusion mechanisms and extended coverage to different physico-chemical environments [49]. Interaction experiments confirmed that consortia composed of *B. velezensis* and *Pediococcus spp.* exhibited antagonistic scores against pathogens at least as high as the best-performing individual strain. Additionally, consortia of *B. velezensis* and *Pediococcus* spp. demonstrated a systematic additive effect against *S. enterica* in the recruitment model, where the positive biofilm is pre-established before pathogen arrival. This effect is not only attributed to pH reduction by *Pediococcus* spp., but rather to spatial and nutritional competition, as supported by our modelling experiments. This finding is consistent with the high surface coverage capacity and improved biofilm production observed in consortia of *B. velezensis* and *Pediococcus* spp.

The use of consortia composed of a *B. velezensis* strain from the Kin consortium and a *Pediococcus* spp. strain results in more intense exclusion of *S. enterica* after 24 hours in the recruitment model compared to the performance of the individual strains. Kinetic experiments over 12 hours in the recruitment model demonstrate that the Kin consortium, when paired with the *P. pentosaceus* R1094 strain, significantly reduces the growth potential of *S. enterica*. This suggests that the combination of the Kin consortium with the R1094 strain allows for more effective early exclusion, which levels out over time with the *P. acidilactici* R1001. Further studies should investigate potential beneficial metabolic exchanges between *B. velezensis* and *Pediococcus* spp. as well as the dynamics of pathogen exclusion during interference. This should be achieved through the use of omics approaches [53] and genome-based modelling [54].

In conclusion, our study highlights the compatibility between selected *B. velezensis* and *Pediococcus* spp. strains, resulting in improved pathogen elimination with a specific additive effect on *S. enterica*. The compatibility and exclusion capacity of strains are intricately linked to biofilm formation, emphasising the importance of pre-establishing positive biofilms on surfaces to achieve additive effects. The imaging methodology presented provides a rational framework for assembling SynComs in a biofilm context, with a focus on strain compatibility and antagonistic potential against pathogens. Moreover, the techniques developed in our study have broad potential applications in biotechnological fields aimed at pathogen control, bioremediation, biopreservation and probiotics.

## Data availability

Raw stacks of images dataset corresponding to the figure 1 and 2 has been deposited in Data INRAE (accession number https://doi.org/10.57745/XRXQEI). Further inquiries can be directed to the corresponding author. Dataset kinetics and R codes used to assess growth parameters are available on Github: https://github.com/lguillier/dataset_kin

## Funding

This study was funded by INRAE, Lallemand SAS, the French National Association for Research and Technology (contracts 2020/0548 and 2024/0397), and the French Agency for Research (ANR LabCom “Biofilm1Health” contract 2024-v1). The funding sources supported the work of all authors except L.G., who did not receive funding from these sources.

## Conflict of interest

VG, CB, GJ, JP-G and MC are employees of the Lallemand SAS company. All the authors declare no competing interests.

## Acknowledgments

We thank the MIMA2 platform (Microscopie et Imagerie des Microorganismes, Animaux et Aliments, https://doi.org/10.15454/1.5572348210007727E12) for microscopic observations. Thanks to Julien Deschamps for the Leica SP8 HCS-CLSM training and Pierre Adenot for the Zeiss LSM 700 observations. Thanks to Harold Guéneau for the Python programming that enabled us to sort the data generated by BiofilmQ. Some figures were created with BioRender (https://www.biorender.com/).

## Author contributions

VG, LG, MC and RB: conceptualization and methodology. MC and RB: validation and supervision. VG, LG, CB: formal analysis and data curation. VG, LG, JP-G, MC, and RB: investigation. PS et M- FN-G contribute to develop the transformation protocol of the strains. GJ generates the phylogenetic tree. MC and RB: resources, project administration, and funding acquisition. VG: writing the original draft. M-FN-G, PS, LG, MC and RB: reviewing, and editing. All authors have read and agreed to the published version of the manuscript.

## Notes

https://doi.org/10.57745/XRXQEI

### Competing Interest Statement

The authors have declared no competing interest.

## References

1. Flemming H-C, Wuertz S. Bacteria and archaea on Earth and their abundance in biofilms. Nat Rev Microbiol 2019;17:247–60.

2. Sauer K, Stoodley P, Goeres DM et al. The biofilm life cycle: expanding the conceptual model of biofilm formation. Nat Rev Microbiol 2022;20:608–20.

3. Flemming H-C, Wingender J, Szewzyk U et al. Biofilms: an emergent form of bacterial life. Nat Rev Microbiol 2016;14:563–75.

4. Flemming H-C, van Hullebusch ED, Neu TR et al. The biofilm matrix: multitasking in a shared space. Nat Rev Microbiol 2023;21:70–86.

5. Dergham Y, Le Coq D, Nicolas P et al. Direct comparison of spatial transcriptional heterogeneity across diverse Bacillus subtilis biofilm communities. Nat Commun 2023;14:7546.

6. Ghoul M, Mitri S. The Ecology and Evolution of Microbial Competition. Trends Microbiol 2016;24:833–45.

7. Palmer JD, Foster KR. Bacterial species rarely work together. Science 2022;376:581–2.

8. Guéneau V, Plateau-Gonthier J, Arnaud L et al. Positive biofilms to guide surface microbial ecology in livestock buildings. Biofilm 2022;4:100075.

9. De Roy K, Marzorati M, Van den Abbeele P et al. Synthetic microbial ecosystems: an exciting tool to understand and apply microbial communities. Environ Microbiol 2014;16:1472–81.

10. van Leeuwen PT, Brul S, Zhang J et al. Synthetic microbial communities (SynComs) of the human gut: design, assembly, and applications. FEMS Microbiol Rev 2023;47:fuad012.

11. Borges F, Briandet R, Callon C et al. Contribution of omics to biopreservation: Toward food microbiome engineering. Front Microbiol 2022;13:951182.

12. Trivedi P, Leach JE, Tringe SG et al. Plant-microbiome interactions: from community assembly to plant health. Nat Rev Microbiol 2020;18:607–21.

13. Heumann A, Assifaoui A, Da Silva Barreira D et al. Intestinal release of biofilm-like microcolonies encased in calcium-pectinate beads increases probiotic properties of Lacticaseibacillus paracasei. NPJ Biofilms Microbiomes 2020;6:44.

14. Guéneau V, Rodiles A, Frayssinet B et al. Positive biofilms to control surface-associated microbial communities in a broiler chicken production system - a field study. Front Microbiol 2022;13:981747.

15. Granato ET, Meiller-Legrand TA, Foster KR. The Evolution and Ecology of Bacterial Warfare. Curr Biol 2019;29:R521–37.

16. Guillier L, Stahl V, Hezard B et al. Modelling the competitive growth between Listeria monocytogenes and biofilm microflora of smear cheese wooden shelves. Int J Food Microbiol 2008;128:51–7.

17. Habimana O, Guillier L, Kulakauskas S et al. Spatial competition with Lactococcus lactis in mixed-species continuous-flow biofilms inhibits Listeria monocytogenes growth. Biofouling 2011;27:1065–72.

18. Maha Swetha BR, Saravanan M, Piruthivraj P. Emerging trends in the inhibition of bacterial molecular communication: An overview. Microb Pathog 2024;186:106495.

19. Shi Y, Wen T, Zhao F et al. Bacteriostasis of nisin against planktonic and biofilm bacteria: Its mechanism and application. J Food Sci 2024;89:1894–916.

20. Byun H, Brockett MR, Pu Q et al. An Intestinal Bacillus velezensis Isolate Displays Broad- Spectrum Antibacterial Activity and Prevents Infection of Both Gram-Positive and Gram-Negative Pathogens In Vivo. J Bacteriol 2023;205:e0013323.

21. Srinivasan S, Jnana A, Murali TS. Modeling Microbial Community Networks: Methods and Tools for Studying Microbial Interactions. Microb Ecol 2024;87:56.

22. Dedrick S, Warrier V, Lemon KP et al. When does a Lotka-Volterra model represent microbial interactions? Insights from in vitro nasal bacterial communities. mSystems 2023;8:e0075722.

23. Pandin C, Le Coq D, Canette A et al. Should the biofilm mode of life be taken into consideration for microbial biocontrol agents? Microb Biotechnol 2017;10:719–34.

24. Guéneau V, Charron R, Costache V et al. Chapter 9 - Spatial analysis of multispecies bacterial biofilms. In: Gurtler V, Patrauchan M (eds.). Methods in Microbiology. Vol 53. Academic Press, 2023, 275–307.

25. Arnaouteli S, Bamford NC, Stanley-Wall NR et al. Bacillus subtilis biofilm formation and social interactions. Nat Rev Microbiol 2021;19:600–14.

26. Mendonça CMN, Oliveira RC, Pizauro LJL et al. Tracking new insights into antifungal and anti- mycotoxigenic properties of a biofilm forming Pediococcus pentosaceus strain isolated from grain silage. Int J Food Microbiol 2023;405:110337.

27. Jang S, Lee D, Jang IS et al. The Culture of Pediococcus pentosaceus T1 Inhibits Listeria Proliferation in Salmon Fillets and Controls Maturation of Kimchi. Food Technol Biotechnol 2015;53:29–37.

28. Zhang N, Wang Z, Shao J et al. Biocontrol mechanisms of Bacillus: Improving the efficiency of green agriculture. Microb Biotechnol 2023;16:2250–63.

29. Olson RD, Assaf R, Brettin T et al. Introducing the Bacterial and Viral Bioinformatics Resource Center (BV-BRC): a resource combining PATRIC, IRD and ViPR. Nucleic Acids Res 2023;51:D678–89.

30. Froger A, Hall JE. Transformation of plasmid DNA into E. coli using the heat shock method. J Vis Exp 2007:253.

31. Dunny GM, Lee LN, LeBlanc DJ. Improved electroporation and cloning vector system for gram- positive bacteria. Appl Environ Microbiol 1991;57:1194–201.

32. Serrano E, Carrasco B, Gilmore JL et al. RecA Regulation by RecU and DprA During Bacillus subtilis Natural Plasmid Transformation. Front Microbiol 2018;9:1514.

33. Dergham Y, Sanchez-Vizuete P, Le Coq D et al. Comparison of the Genetic Features Involved in Bacillus subtilis Biofilm Formation Using Multi-Culturing Approaches. Microorganisms 2021;9, DOI: 10.3390/microorganisms9030633.

34. Pogliano J, Osborne N, Sharp MD et al. A vital stain for studying membrane dynamics in bacteria: a novel mechanism controlling septation during Bacillus subtilis sporulation. Mol Microbiol 1999;31:1149–59.

35. Hartmann R, Jeckel H, Jelli E et al. Quantitative image analysis of microbial communities with BiofilmQ. Nat Microbiol 2021;6:151–6.

36. Cornu M, Billoir E, Bergis H et al. Modeling microbial competition in food: Application to the behavior of Listeria monocytogenes and lactic acid flora in pork meat products. Food Microbiology 2011;28:639–47.

37. Ross T, Dalgaard P, Tienungoon S. Predictive modelling of the growth and survival of Listeria in fishery products. International Journal of Food Microbiology 2000;62:231–45.

38. Baty F, Delignette-Muller M-L, Siberchicot A. nlsMicrobio: Nonlinear Regression in Predictive Microbiology. 2024.

39. Mühlbauer LK, Schulze M, Harpole WS et al. gauseR: Simple methods for fitting Lotka-Volterra models describing Gause’s “Struggle for Existence.” Ecol Evol 2020;10:13275–83.

40. Li Q, Lei Y, Li T. DNA metabarcoding reveals ecological patterns and driving mechanisms of archaeal, bacterial, and eukaryotic communities in sediments of the Sansha Yongle Blue Hole. Sci Rep 2024;14:6745.

41. Song H-S, Lee N-R, Kessell AK et al. Kinetics-based inference of environment-dependent microbial interactions and their dynamic variation. mSystems 2024;9:e0130523.

42. Kimelman H, Shemesh M. Probiotic Bifunctionality of Bacillus subtilis-Rescuing Lactic Acid Bacteria from Desiccation and Antagonizing Pathogenic Staphylococcus aureus. Microorganisms 2019;7:407.

43. Widder S, Allen RJ, Pfeiffer T et al. Challenges in microbial ecology: building predictive understanding of community function and dynamics. ISME J 2016;10:2557–68.

44. Wucher BR, Winans JB, Elsayed M et al. Breakdown of clonal cooperative architecture in multispecies biofilms and the spatial ecology of predation. Proc Natl Acad Sci U S A 2023;120:e2212650120.

45. Chang C-Y, Bajic D, Vila J et al. Emergent coexistence in multispecies microbial communities. 2022:2022.05.20.492860.

46. Lyng M, Jørgensen JPB, Schostag MD et al. Competition for iron shapes metabolic antagonism between Bacillus subtilis and Pseudomonas marginalis. ISME J 2024;18:wrad001.

47. Keshmirshekan A, de Souza Mesquita LM, Ventura SPM. Biocontrol manufacturing and agricultural applications of Bacillus velezensis. Trends Biotechnol 2024;42:986–1001.

48. Pandin C, Darsonval M, Mayeur C et al. Biofilm Formation and Synthesis of Antimicrobial Compounds by the Biocontrol Agent Bacillus velezensis QST713 in an Agaricus bisporus Compost Micromodel. Appl Environ Microbiol 2019;85:e00327–19.

49. Spragge F, Bakkeren E, Jahn MT et al. Microbiome diversity protects against pathogens by nutrient blocking. Science 2023;382:eadj3502.

50. Stefanic P, Kraigher B, Lyons NA et al. Kin discrimination between sympatric Bacillus subtilis isolates. Proc Natl Acad Sci U S A 2015;112:14042–7.

51. Kraigher B, Butolen M, Stefanic P et al. Kin discrimination drives territorial exclusion during Bacillus subtilis swarming and restrains exploitation of surfactin. ISME J 2022;16:833–41.

52. Liu Y, Huang R, Chen Y et al. Involvement of Flagellin in Kin Recognition between Bacillus velezensis Strains. mSystems 2022;7:e0077822.

53. Delavy M, Sertour N, d’Enfert C et al. Metagenomics and metabolomics approaches in the study of Candida albicans colonization of host niches: a framework for finding microbiome-based antifungal strategies. Trends Microbiol 2023;31:1276–86.

54. Wu S, Qu Z, Chen D et al. Deciphering and designing microbial communities by genome-scale metabolic modelling. Comput Struct Biotechnol J 2024;23:1990–2000.

