## Supplementary materials for "Guided assembly of multispecies positive biofilms targeting undesirable bacteria"

#### Sup.1 Characteristics of strains used in this study.

| Name | Origine and genotype | Reference |
|---|---|---|
| *Bacillus velezensis* 11285 | unknown | Lallemand Aquapharm |
| *Bacillus velezensis* 12048 | unknown | Lallemand Aquapharm |
| *Bacillus velezensis* ILPB8 | Surfaces of commercial broiler chicken houses | [1] |
| *Bacillus velezensis* 12701 | unknown | Lallemand Aquapharm |
| *Bacillus velezensis* 11457 | unknown | Lallemand Aquapharm |
| *Bacillus velezensis* B18 | Surfaces of commercial broiler chicken houses | [1] |
| *Bacillus velezensis* B1 | Surfaces of commercial broiler chicken houses | [1] |
| *Bacillus velezensis* 12832 | unknown | Lallemand Aquapharm |
| *Bacillus velezensis* 1273 | Very wet soil of a snow combe, 2400m altitude | INRAE B3D |
| *Bacillus velezensis* 12001 | unknown | Lallemand Aquapharm |
| *Bacillus subtilis* 1202 | Water from a river | INRAE B3D |
| *Bacillus licheniformis* 1234 | Small greenish pebbles | INRAE B3D |
| *Bacillus licheniformis* C5 | Surfaces of commercial broiler chicken houses | [1] |
| *Bacillus licheniformis* 1218 | Sea urchin intestine in sea water | INRAE B3D |
| *Bacillus licheniformis* 1298 | Pinkish grey gravel | INRAE B3D |
| *Bacillus licheniformis* 1219 | Greyish sandy soil | INRAE B3D |
| *Paenibacillus* sp. 1167 | Black soil | INRAE B3D |
| *Peanibacillus* sp. 1399 | Dark brown soil | INRAE B3D |
| *Pediococcus acidilactici* R1001 | unknown | Lallemand |
| *Pediococcus pentosaceus* R1094 | unknown | Lallemand |
| *Bacillus velezensis* 11285 mCherry | *Bacillus velezensis* 11285 with pGM11-*mcherry* (ery) | This study |
| *Bacillus velezensis* 12048 GFP | *Bacillus velezensis* 12048 with pCM11-*gfp* (ery) | This study |
| *Bacillus velezensis* ILPB8 GFP | *Bacillus velezensis* ILPB8 with pCM11-*gfp* (ery) | This study |
| *Bacillus velezensis* ILPB8 mCherry | *Bacillus velezensis* ILPB8 with pGM11-*mcherry* (ery) | This study |
| *Staphylococcus aureus* RN4220 | Laboratory strain with pALC2084-*gfp* (ery) | [2] |
| *Enterococcus cecorum* DSM20682 | Strain from DSM collection, isolated from chicken caecum with pCM11-*gfp* (ery) | This study |
| *Salmonella enterica enterica* serotype *enteritidis* NCTC6676 | Strain from NCTC collection, isolated from a dead cow with pCM11-*gfp* (amp) | This study |
| *Escherichia coli* CIRMBP0248 | Strain from CIRMBP collection, isolated from chicken intestine with pCM11-*gfp* (amp) | This study |

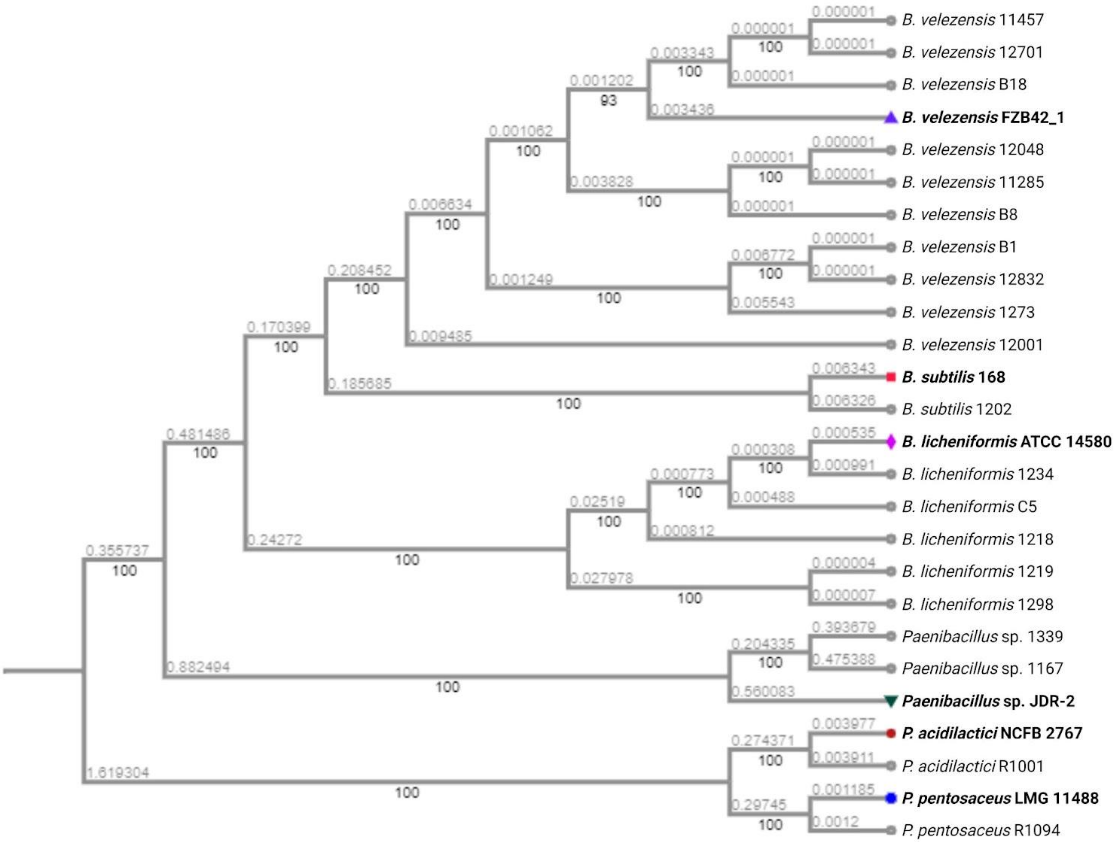


**Sup. 2 Phylogenetic tree of the 20 candidate beneficial strains used in the study along with reference strains.** The tree was obtained using the Bacterial Phylogenetic Tree Service of BV-BRC [3]**.** Reference strains have been added to the tree in bold.

#### Supplementary Methods

##### Sup 3: Bacterial transformation protocols:

Various protocols were used to transform plasmids pCM11 and pGM11 into strains. These plasmids are constitutively expressing a fluorophore (Gfp or mCherry) for the purpose of conducting CLSM studies.

##### Chemical transformation of *E. coli* and *S. enterica*

Briefly, competent cells were prepared by adding 1 ml of overnight culture of the wild-type strain to 80 ml of LB (Difco™) and incubated at 37°C until an OD600 of 0.3-0.4 was reached. The culture was placed on ice for 15 min, split in two 50 mL tubes and then centrifuged at 6000 g for 10 minutes. The supernatant was removed and the pellets were resuspended in 30 mL of 0.1 M CaCl2. After a 30-minute incubation at 30 °C, the culture was centrifuged again at 6000 g for 10 min. The supernatant was removed and the pellets were suspended in 3 mL of 0.1M Cacl2 with 15 % glycerol. Aliquots of 50 µL were prepared and stored at -80°C.

For transformation, a tube of competent cells was placed on ice in order to defrost gently. 1 µg of pCM11 plasmid was then added in the tube, and incubated on ice for 15 minutes. The mixture was heat-shocked at 42 °C for 90 seconds and then placed back on ice for a few seconds. A volume of 250 µL of SOC medium was then added, and the tube was incubated at 37 °C for 1 hour with agitation. The culture was plate on LB agar supplemented with 100 µg/mL ampicillin and incubated overnight at 37 °C. The resulting clones that carried the plasmid were stored.

##### Electroporation of *E. cecorum*

*E. cecorum* transformation was performed by electroporation according to an already established protocol [4]. To obtain electrocompetent cells, SGM17 medium (37,25 g M17 Broth from Difco™; 5 g/L glucose; 0.16 M saccharose; 2 % glycine) was inoculated 1:100 (v;v) with an overnight culture of wild type *E. cecorum* and incubated under agitation at 37°C until reaching an OD600 of 0.5. The culture was placed on ice for 10 min. All subsequent steps were performed on ice. The bacteria were centrifuged at 3800 g for 10 min at 4°C, and the supernatant was removed. The pellet was resuspended in the same volume of ice-cold Sac-Gly solution (saccharose 0.5 M, glycerol 10%, pH 7.0). This washing step was repeated 5 times, dividing each time per 2 the volume of Sac-Gly to suspend the pellet. The final wash consists in resuspending the pellet in Sac-Gly to 1/150e of the initial culture volume. Aliquots of 40 µL were prepared, frozen in liquid nitrogen and stored at -80°C.

For transformation, 40 µL of competent cells were defrosted on ice and 1 µg of pCM11 plasmid was added. The mixture was placed in a 2 mm wide electroporation cuvette and the electric pulse was applied (25 µF, 200 Ohms, 2.4 kV). Immediately, 1 mL of SGM17 medium supplemented with 0.5 M MgCl2 and 10 mM CaCl2 was added and the tube placed on ice for 5 min. The mixture was placed at 30°C without agitation for 3h. The culture was plated on M17 supplemented with 0.5 g/L of glucose and erythromycin at 1 µg/mL and incubated 2 days at 30 °C. The resulting clones that carried the plasmid were stored.

**Transformation of naturally competent *B. velezensis***

Before starting, 5 µg of plasmid (pCM11 or pGM11) was linearised by the EcoRI restriction enzyme (NEB), and the enzyme was subsequently inactivated. This step was followed by a ligation using the Quick ligation kit (NEB) to create plasmid concatemers. From a TSA plate, one colony was inoculated into 1 mL of MC medium (20 g/L glucose; 10 g/L potassium glutamate; 1 M pH7 potassium phosphate buffer; 0.1 M sodium trisodium citrate; 10 g/L ferric ammonium citrate; 5 % casein hydrolysate; 1% tryptophan; 1M MgSO4; completed with H2O). The culture was incubated for 4 h at 37°C, 180 rpm. 500 µL of the culture was collected and added to a new tube containing the 5 µg of pCM11 concatemers. The mixture was incubated 1.5 hours at 37°C with shaking at 180 rpm. 200 µL of each culture was plated on TSA supplemented with erythromycin 0.5 µg/mL and lincomycin 25 µg/mL. The plates were incubated at 30°C for 2 days. The resulting clones that carried the plasmid were stored.

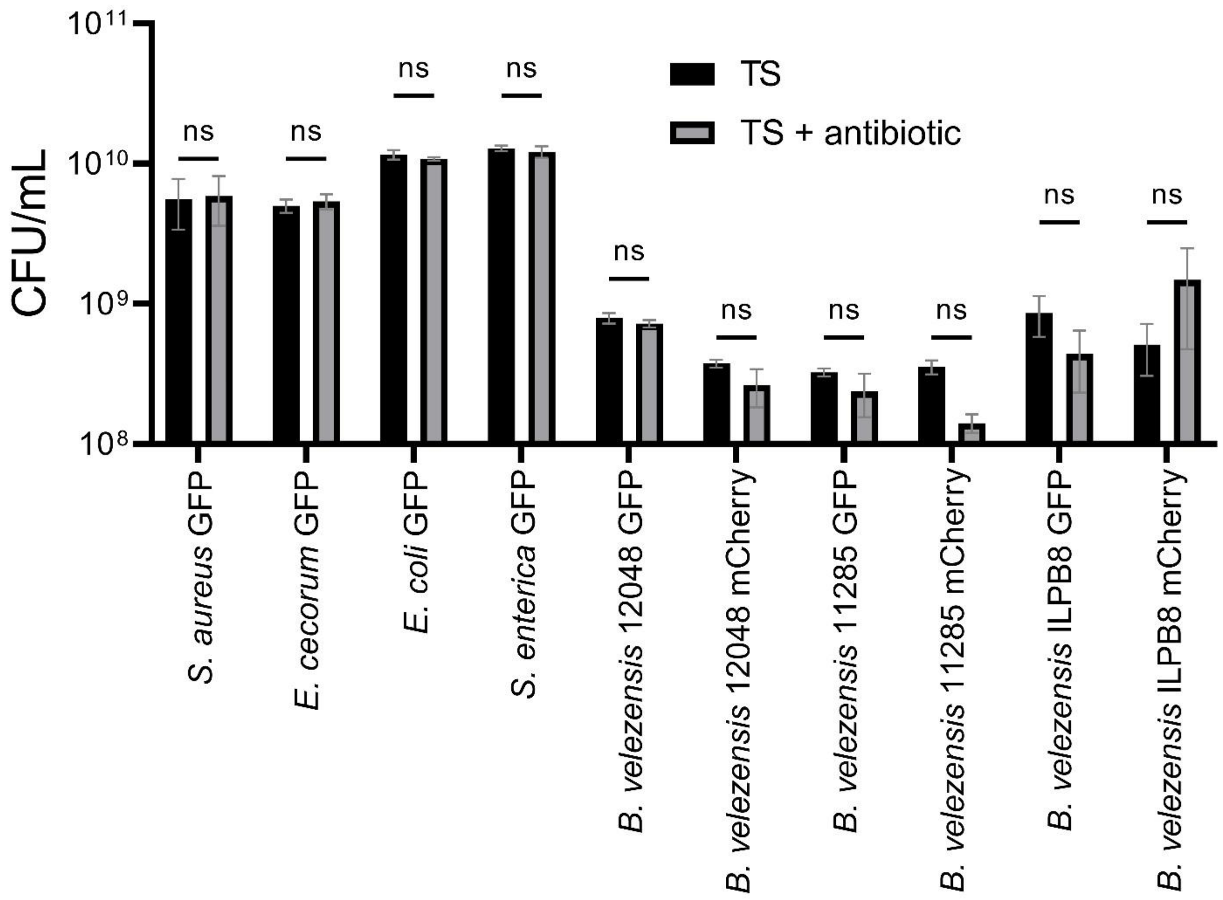


**Sup. 4 Plasmid stability in biofilm experiments.** Biofilms were cultured in 96 wells plate in 200 µL of TSB for 24 hours without antibiotics from overnight cultures of bacteria initially grown in the presence of antibiotics. Post-cultivation, biofilms were detached and homogenised in their related wells. Plasmid stability was assessed by plating on TSA or TSA supplemented with antibiotics (ery 5 µg/mL for *S. aureus* and *E. cecorum* and amp 100 µg/mL for *E. coli* and *S. enterica*). Three biological replicates were performed for each condition. Error bars correspond to standard deviation of the measurements.

71

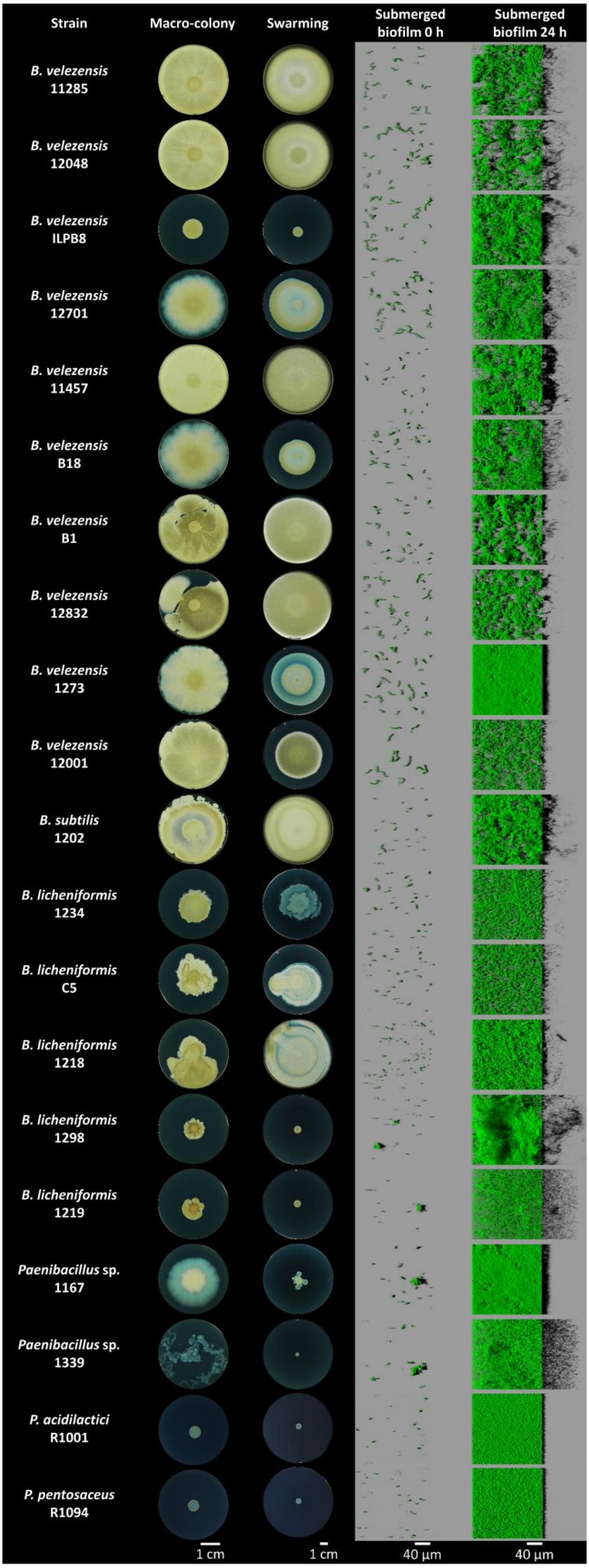

Strain
Macro-colony
Swarming
Submerged biofilm 0 h
Submerged biofilm 24 h
B. velezensis 11285
B. velezensis 12048
B. velezensis ILPB8
B. velezensis 12701
B. velezensis 11457
B. velezensis B18
B. velezensis B1
B. velezensis 12832
B. velezensis 1273
B. velezensis 12001
B. subtilis 1202
B. licheniformis 1234
B. licheniformis C5
B. licheniformis 1218
B. licheniformis 1298
B. licheniformis 1219
Paenibacillus sp. 1167
Paenibacillus sp. 1339
P. acidilactici R1001
P. pentosaceus R1094
1 cm
1 cm
40 µm
40 µm

**Sup. 5 Phenotypic characterisation of *Bacillus* strains in different biofilm models.** To initiate the macro-colony model experiments, 5 ml cultures were prepared in trypticase soy broth (TSB; Biomérieux, France) from glycerol stock stored at -80°C. These cultures were incubated at 30°C overnight without agitation. Following homogenisation by rapid vortexing for 5 seconds, 3 µL of culture were transferred to a well of a six-well plate containing 4 ml of TSA 1.5% agar. The samples were dried under a hood for 10 minutes and incubated at 30°C for 4 days. For the swarming experiment, a similar protocol was followed, but with 20 mL of TSA 0.7% agar in a Petri dish, incubated overnight at 30°C. Mono-species submerged biofilms were observed using the CLSM. Overnight cultures were diluted at 1:100e in TSB, and 200 µL of this dilution was added to each well of a 96-well plate. The cultures were allowed to adhere for 1 hour and 30 minutes. After the adhesion phase, the supernatant was carefully removed and replaced with fresh TSB. The plates were then incubated for 24 hours at 30°C without agitation, and 432 image stacks were collected. Representative images are shown.

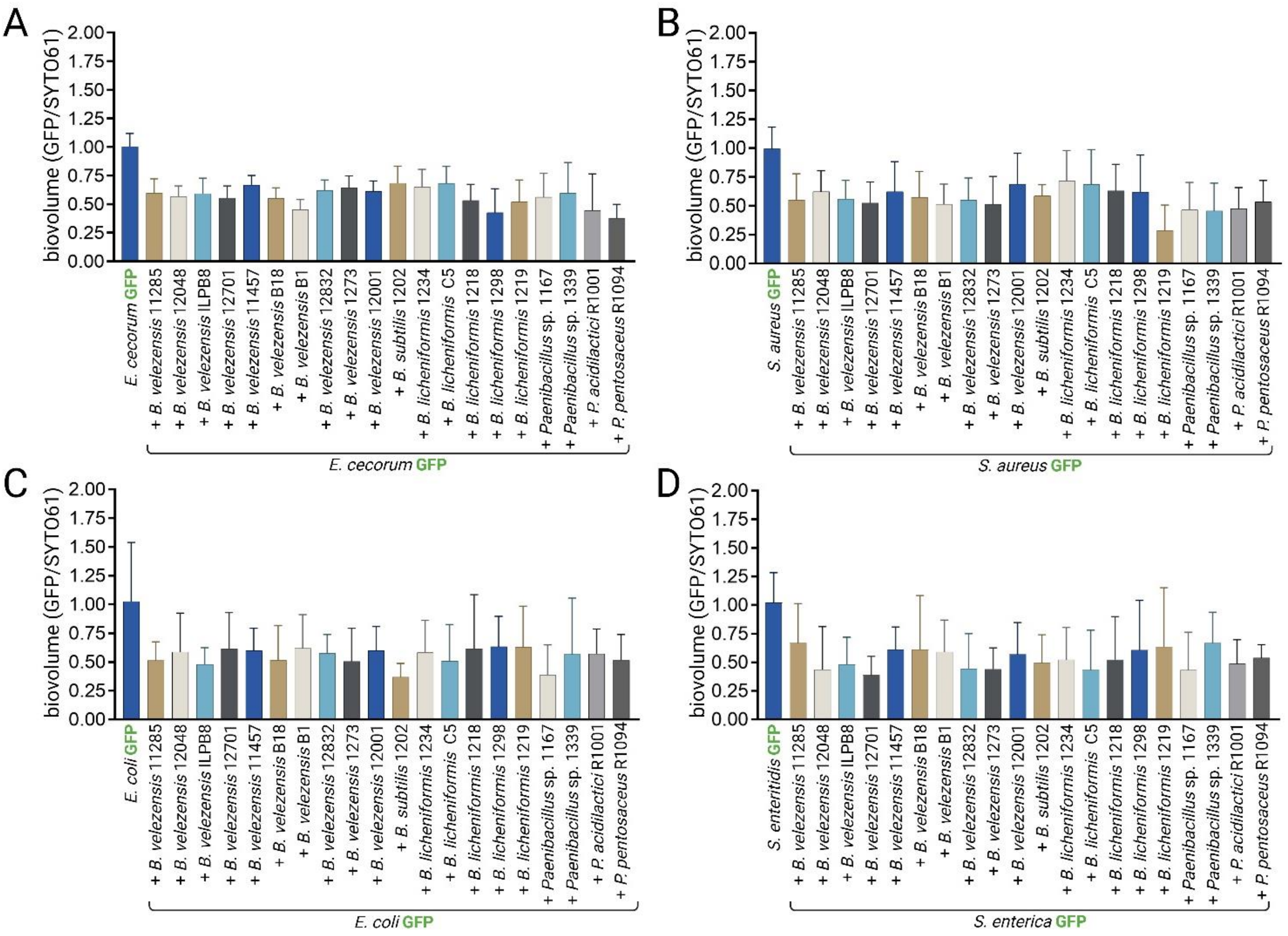


**Sup. 6 Adhesion ratio of candidate beneficial strains and GFP-labelled pathogens in the co-inoculation growth model.** Volumes of overnight cultures of non-labeled beneficial strains were adjusted to achieve an equal biovolume of GFP-labelled pathogens and beneficial strains adhered to the bottom of the wells. Pathogens were genetically marked with GFP, and the entire population was chemically labelled with SYTO61 after 1 h 30 min of adhesion. A GFP biovolume to SYTO61 biovolume ratio of 0.5 indicates an equal quantity of *B. velezensis* 12048 GFP and beneficial strain.

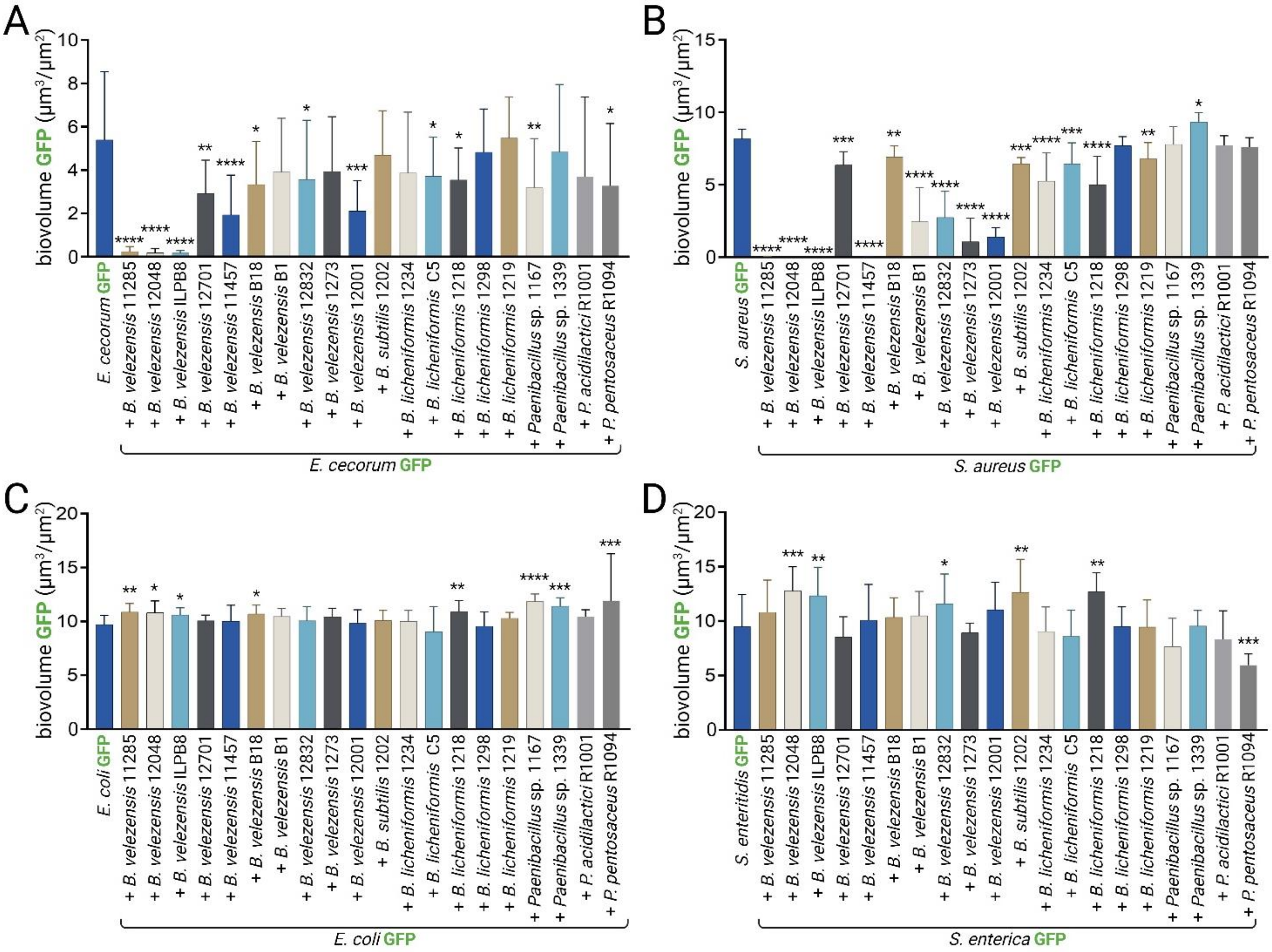


**Sup. 7 Biovolume of GFP-labelled pathogens in the co-inoculation growth model with candidate beneficial strains.** The biovolume of GFP-labelled pathogens co-cultured with candidate beneficial strains was measured. The results are shown for (A) GFP-labelled *E. cecorum*, (B) GFP-labelled *S. aureus*, (C) GFP-labelled *E. coli*, and (D) GFP-labelled *S. enterica*. Error bars correspond to standard deviation. The biovolume of the GFP-labelled pathogen in the presence of the candidate beneficial strain was compared to the biovolume of the pathogen grown alone.

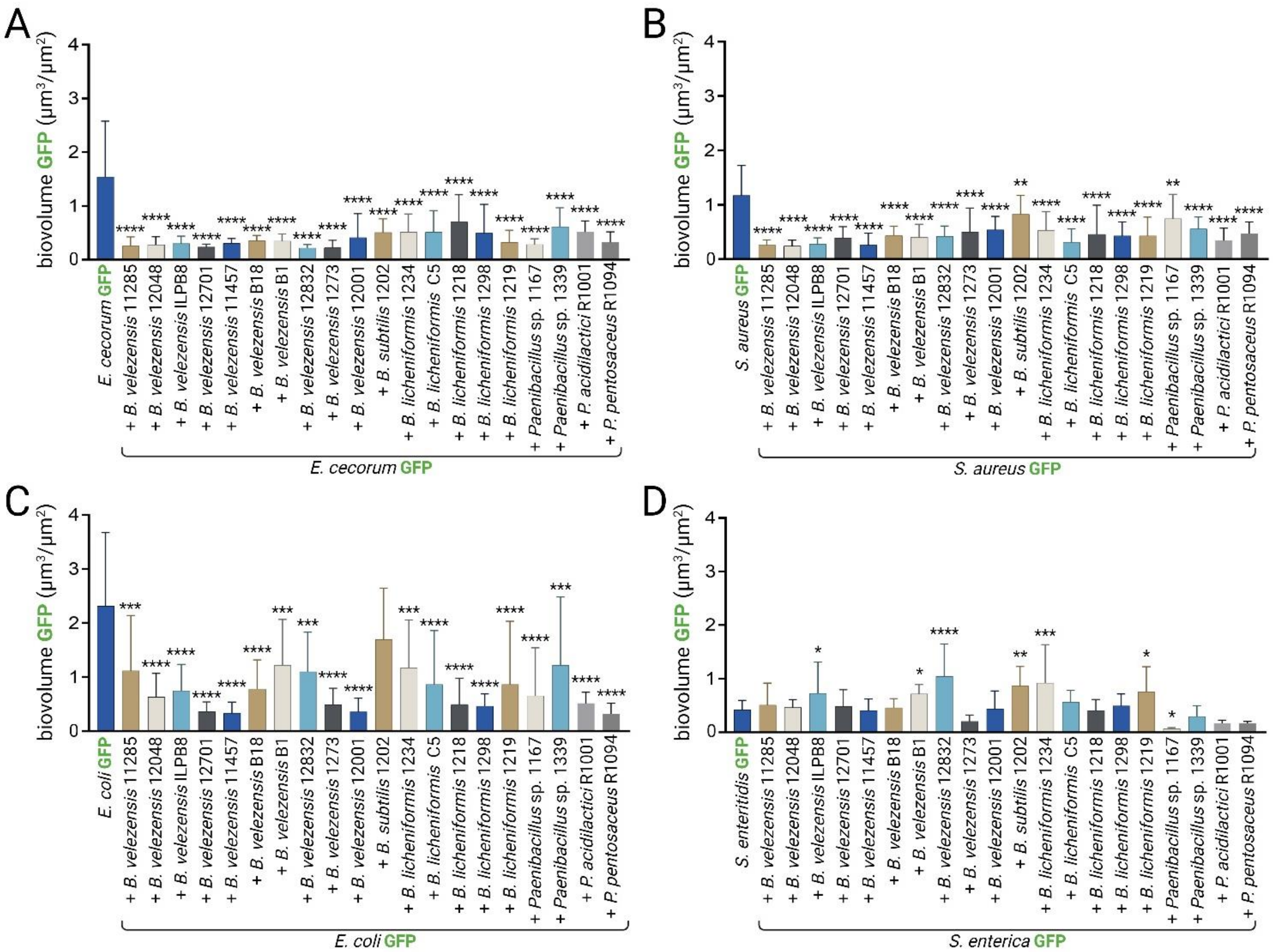


**Sup. 8 Biovolume of GFP-labelled pathogen in the recruitment t=0 h growth model co-cultured with candidate beneficial strains.** The biovolume of GFP-labelled pathogens in the recruitment t=0 h growth model, co-cultured with candidate beneficial strains, was measured. The results are shown for (A) GFP-labelled *E. cecorum*, (B) GFP-labelled *S. aureus*, (C) GFP-labelled *E. coli*, and (D) GFP-labelled *S. enterica*. Error bars correspond to standard deviation. The biovolume of the GFP-labelled pathogen in the presence of the candidate beneficial strains was compared to the biovolume of the pathogen grown alone.

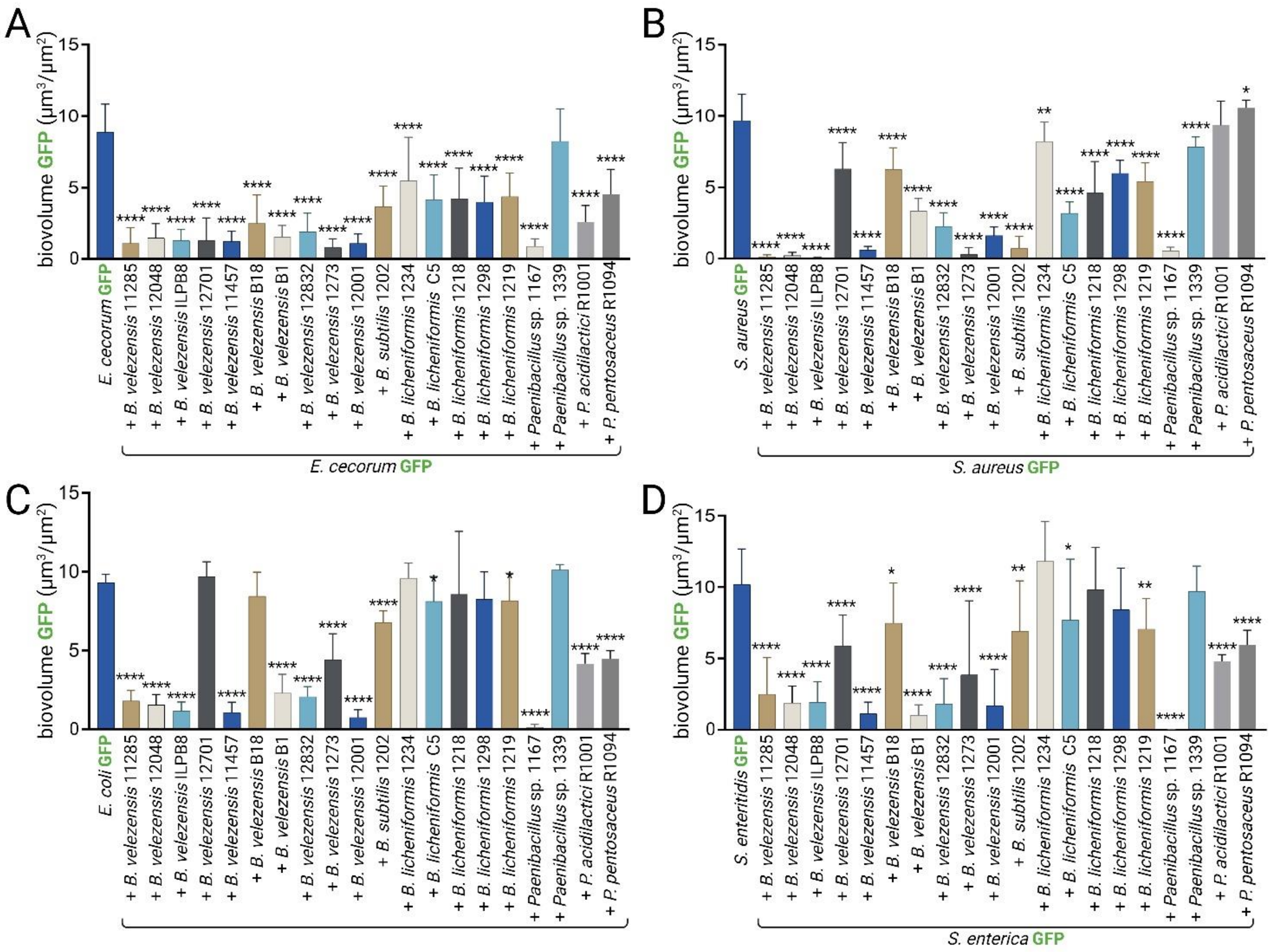


**Sup. 9 Biovolume of GFP-labelled pathogen in the recruitment t=24 hours growth model co-cultured with candidate beneficial strains.** The results are shown for (A) GFP-labelled *E. cecorum*, (B) GFP-labelled *S. aureus*, (C) GFP-labelled *E. coli*, and (D) GFP-labelled *S. enterica*. Error bars correspond to standard deviation. The biovolume values of the GFP-labelled pathogen in the presence of the candidate beneficial strains were compared with the biovolume of the pathogen grown alone.

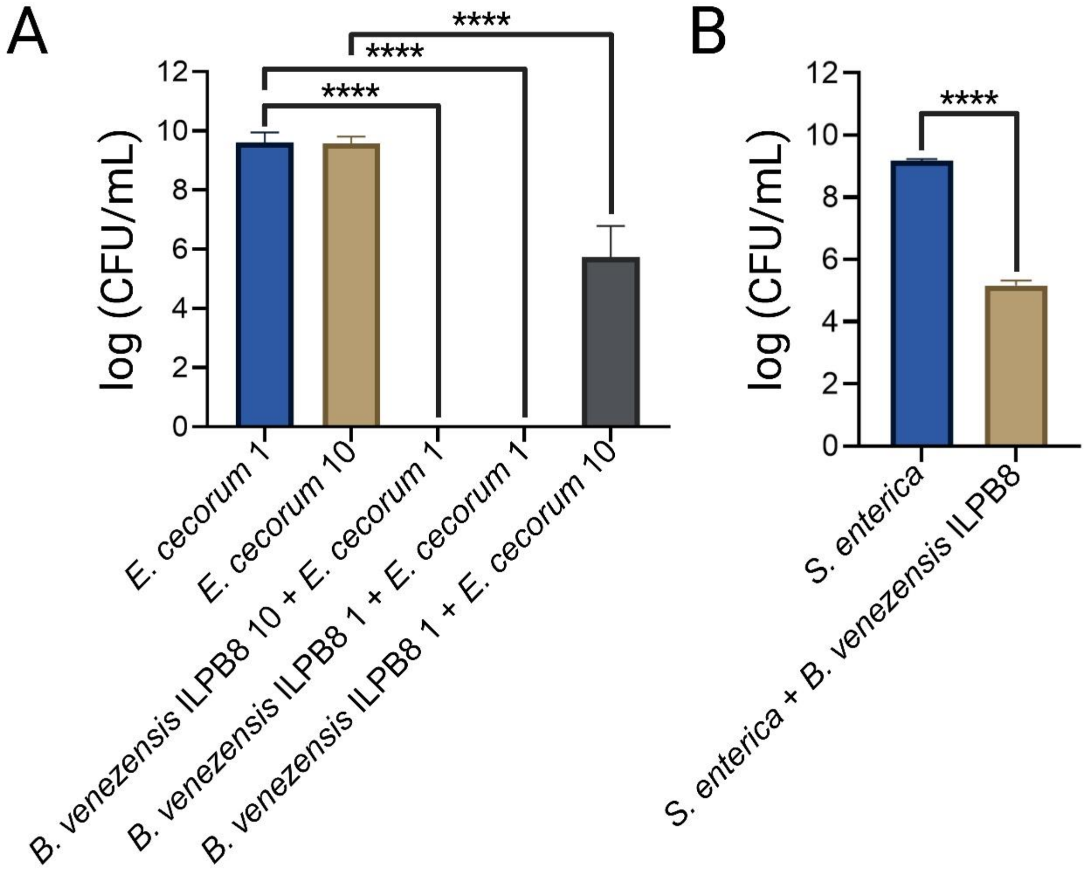


**Sup. 10 Enumeration of pathogens after 24 hours of incubation in the presence or absence of *B. velezensis* ILPB8.** Pathogen counts were measured in submerged biofilms after 24 hours of incubation with and without *B. velezensis* ILPB8. Biofilms were vortexed 10 s vigorously before enumeration. (A) Results with *E. cecorum* using different ratios of the two partners at the start of the co-inoculation experiment. (B) Results with *S. enterica* in the recruitment model. Experiments were performed with 3 biological replicates. Error bars correspond to standard deviation.

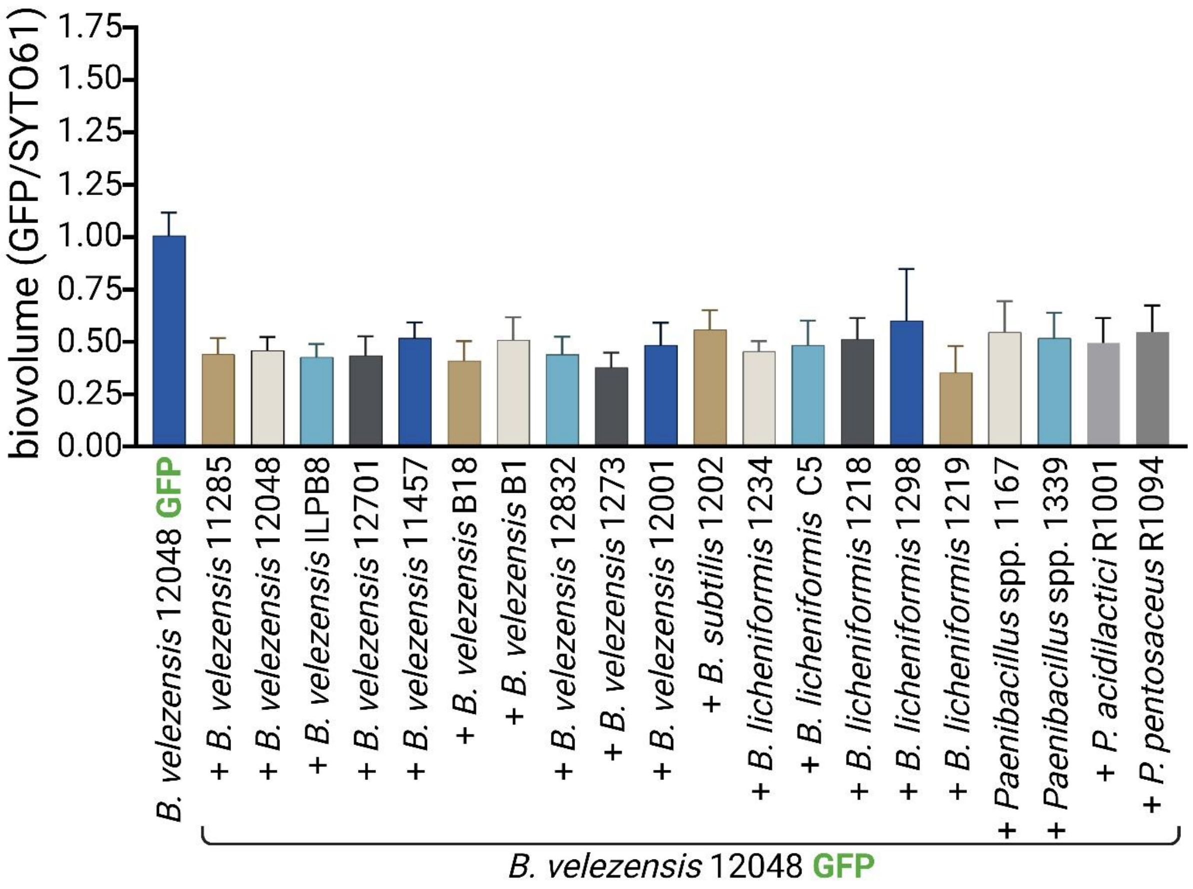


**Sup. 11 Adhesion ratio of the 20 beneficial strains and GFP-labeled *B. velezensis* in the co-inoculation growth model.** (A) Volumes from overnight cultures were calibrated to achieve an equal biovolume of *B. velezensis* GFP and non-labeled *Bacillus* spp. strains adhered to the bottom of the wells. *B. velezensis* 12048 was genetically marked with GFP, and the entire population was chemically labelled with SYTO61 after 1 h 30 min of adhesion. A GFP biovolume to SYTO61 biovolume ratio of 0.5 indicates an equal quantity of *B. velezensis* 12048 GFP and beneficial strains. (B) Verification of the adhesion ratio was performed with GFP-labelled *B. velezensis* 11285, 12048 and ILPB8 and *Pediococcus* spp. Error bars correspond to standard deviation.

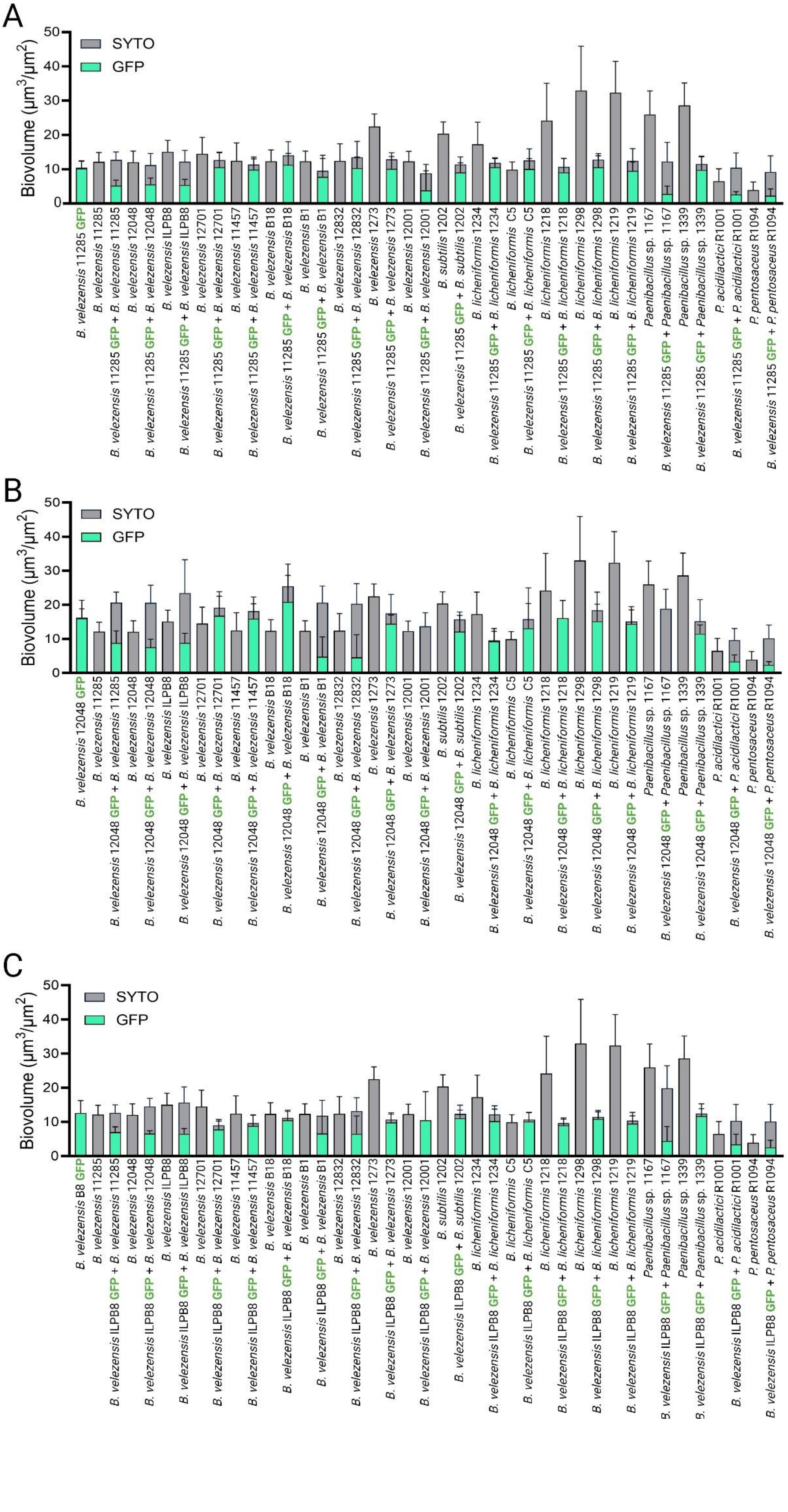
A
Biovolume (µm³/µm²)
0
10
20
30
40
50
SYTO
GFP
B. velezensis 11285 GFP
B. velezensis 11285
B. velezensis 11285 GFP + B. velezensis 11285
B. velezensis 12048
B. velezensis 11285 GFP + B. velezensis 12048
B. velezensis ILPB8
B. velezensis 11285 GFP + B. velezensis ILPB8
B. velezensis 12701
B. velezensis 11285 GFP + B. velezensis 12701
B. velezensis 11457
B. velezensis 11285 GFP + B. velezensis 11457
B. velezensis B18
B. velezensis 11285 GFP + B. velezensis B18
B. velezensis B1
B. velezensis 11285 GFP + B. velezensis B1
B. velezensis 12832
B. velezensis 11285 GFP + B. velezensis 12832
B. velezensis 1273
B. velezensis 11285 GFP + B. velezensis 1273
B. velezensis 12001
B. velezensis 11285 GFP + B. velezensis 12001
B. subtilis 1202
B. velezensis 11285 GFP + B. subtilis 1202
B. licheniformis 1234
B. velezensis 11285 GFP + B. licheniformis 1234
B. licheniformis C5
B. velezensis 11285 GFP + B. licheniformis C5
B. licheniformis 1218
B. velezensis 11285 GFP + B. licheniformis 1218
B. licheniformis 1298
B. velezensis 11285 GFP + B. licheniformis 1298
B. licheniformis 1219
B. velezensis 11285 GFP + B. licheniformis 1219
Paenibacillus sp. 1167
B. velezensis 11285 GFP + Paenibacillus sp. 1167
Paenibacillus sp. 1339
B. velezensis 11285 GFP + Paenibacillus sp. 1339
P. acidilactici R1001
B. velezensis 11285 GFP + P. acidilactici R1001
P. pentosaceus R1094
B. velezensis 11285 GFP + P. pentosaceus R1094
B
Biovolume (µm³/µm²)
0
10
20
30
40
50
SYTO
GFP
B. velezensis 12048 GFP
B. velezensis 11285
B. velezensis 12048 GFP + B. velezensis 11285
B. velezensis 12048
B. velezensis 12048 GFP + B. velezensis 12048
B. velezensis ILPB8
B. velezensis 12048 GFP + B. velezensis ILPB8
B. velezensis 12701
B. velezensis 12048 GFP + B. velezensis 12701
B. velezensis 11457
B. velezensis 12048 GFP + B. velezensis 11457
B. velezensis B18
B. velezensis 12048 GFP + B. velezensis B18
B. velezensis B1
B. velezensis 12048 GFP + B. velezensis B1
B. velezensis 12832
B. velezensis 12048 GFP + B. velezensis 12832
B. velezensis 1273
B. velezensis 12048 GFP + B. velezensis 1273
B. velezensis 12001
B. velezensis 12048 GFP + B. velezensis 12001
B. subtilis 1202
B. velezensis 12048 GFP + B. subtilis 1202
B. licheniformis 1234
B. velezensis 12048 GFP + B. licheniformis 1234
B. licheniformis C5
B. velezensis 12048 GFP + B. licheniformis C5
B. licheniformis 1218
B. velezensis 12048 GFP + B. licheniformis 1218
B. licheniformis 1298
B. velezensis 12048 GFP + B. licheniformis 1298
B. licheniformis 1219
B. velezensis 12048 GFP + B. licheniformis 1219
Paenibacillus sp. 1167
B. velezensis 12048 GFP + Paenibacillus sp. 1167
Paenibacillus sp. 1339
B. velezensis 12048 GFP + Paenibacillus sp. 1339
P. acidilactici R1001
B. velezensis 12048 GFP + P. acidilactici R1001
P. pentosaceus R1094
B. velezensis 12048 GFP + P. pentosaceus R1094
C
Biovolume (µm³/µm²)
0
10
20
30
40
50
SYTO
GFP
B. velezensis B8 GFP
B. velezensis 11285
B. velezensis ILPB8 GFP + B. velezensis 11285
B. velezensis 12048
B. velezensis ILPB8 GFP + B. velezensis 12048
B. velezensis ILPB8
B. velezensis ILPB8 GFP + B. velezensis ILPB8
B. velezensis 12701
B. velezensis ILPB8 GFP + B. velezensis 12701
B. velezensis 11457
B. velezensis ILPB8 GFP + B. velezensis 11457
B. velezensis B18
B. velezensis ILPB8 GFP + B. velezensis B18
B. velezensis B1
B. velezensis ILPB8 GFP + B. velezensis B1
B. velezensis 12832
B. velezensis ILPB8 GFP + B. velezensis 12832
B. velezensis 1273
B. velezensis ILPB8 GFP + B. velezensis 1273
B. velezensis 12001
B. velezensis ILPB8 GFP + B. velezensis 12001
B. subtilis 1202
B. velezensis ILPB8 GFP + B. subtilis 1202
B. licheniformis 1234
B. velezensis ILPB8 GFP + B. licheniformis 1234
B. licheniformis C5
B. velezensis ILPB8 GFP + B. licheniformis C5
B. licheniformis 1218
B. velezensis ILPB8 GFP + B. licheniformis 1218
B. licheniformis 1298
B. velezensis ILPB8 GFP + B. licheniformis 1298
B. licheniformis 1219
B. velezensis ILPB8 GFP + B. licheniformis 1219
Paenibacillus sp. 1167
velezensis ILPB8 GFP + Paenibacillus sp. 1167
Paenibacillus sp. 1339
velezensis ILPB8 GFP + Paenibacillus sp. 1339
P. acidilactici R1001
B. velezensis ILPB8 GFP + P. acidilactici R1001
P. pentosaceus R1094
B. velezensis ILPB8 GFP + P. pentosaceus R1094

**Sup. 12 GFP-labelled *B. velezensis* biovolume in the co-inoculation growth model co-cultured with 20 beneficial candidate strains.** The biovolume of GFP-labelled *B. velezensis* co-cultured with 20 beneficial candidate strains was measured. The results are shown for (A) GFP-labelled *B. velezensis* 11285, (B) GFP-labelled *B. velezensis* 12048, and (C) GFP-labelled *B. velezensis* ILPB8. Error bars correspond to standard deviation.

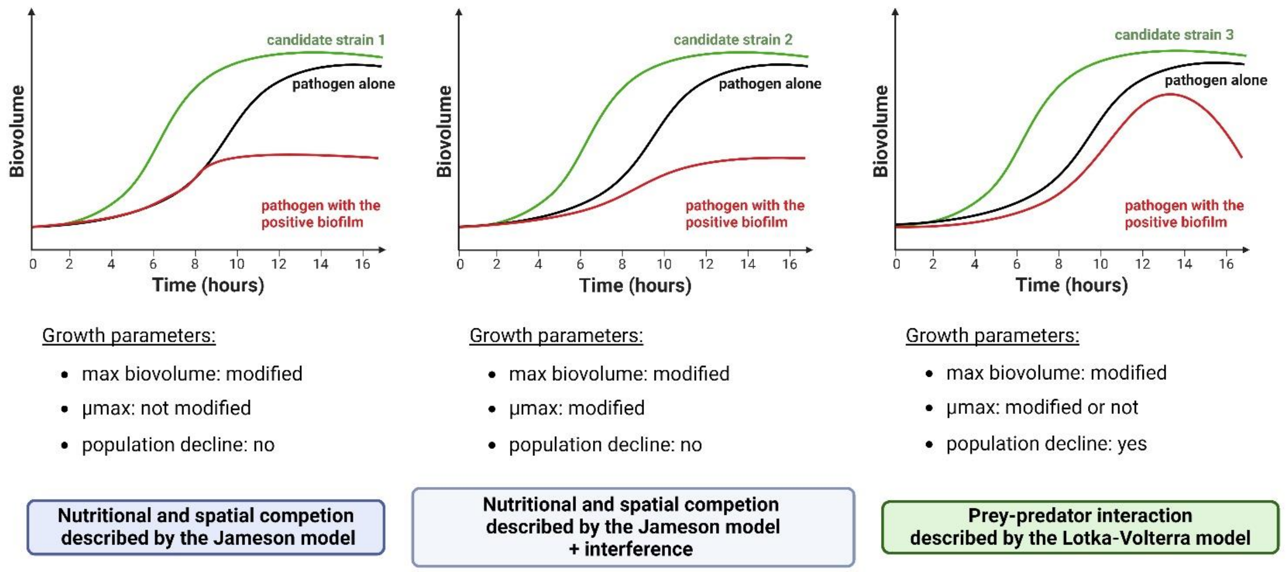


**Sup. 13 Dynamics of biofilm formation through CLSM kinetic studies.** Schematic representation of potential interactions at CLSM and the resulting families of mechanisms.

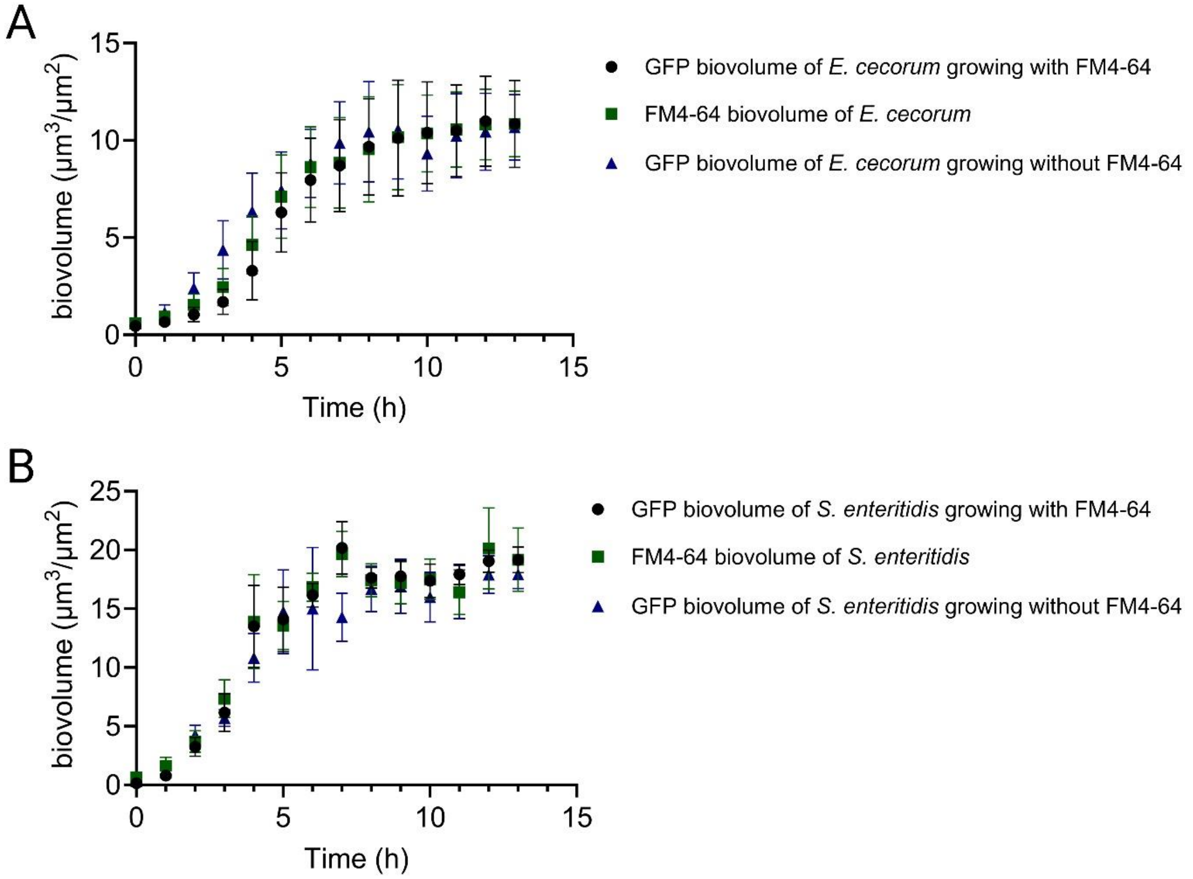


**Sup. 14 Effect of the FM4-64 on pathogen growth.** (A) Biovolumes of *E. cecorum* GFP biofilms grown with or without FM4-64 and (B) Biovolume of *S. enterica* GFP biofilms grown with and without FM4-64.

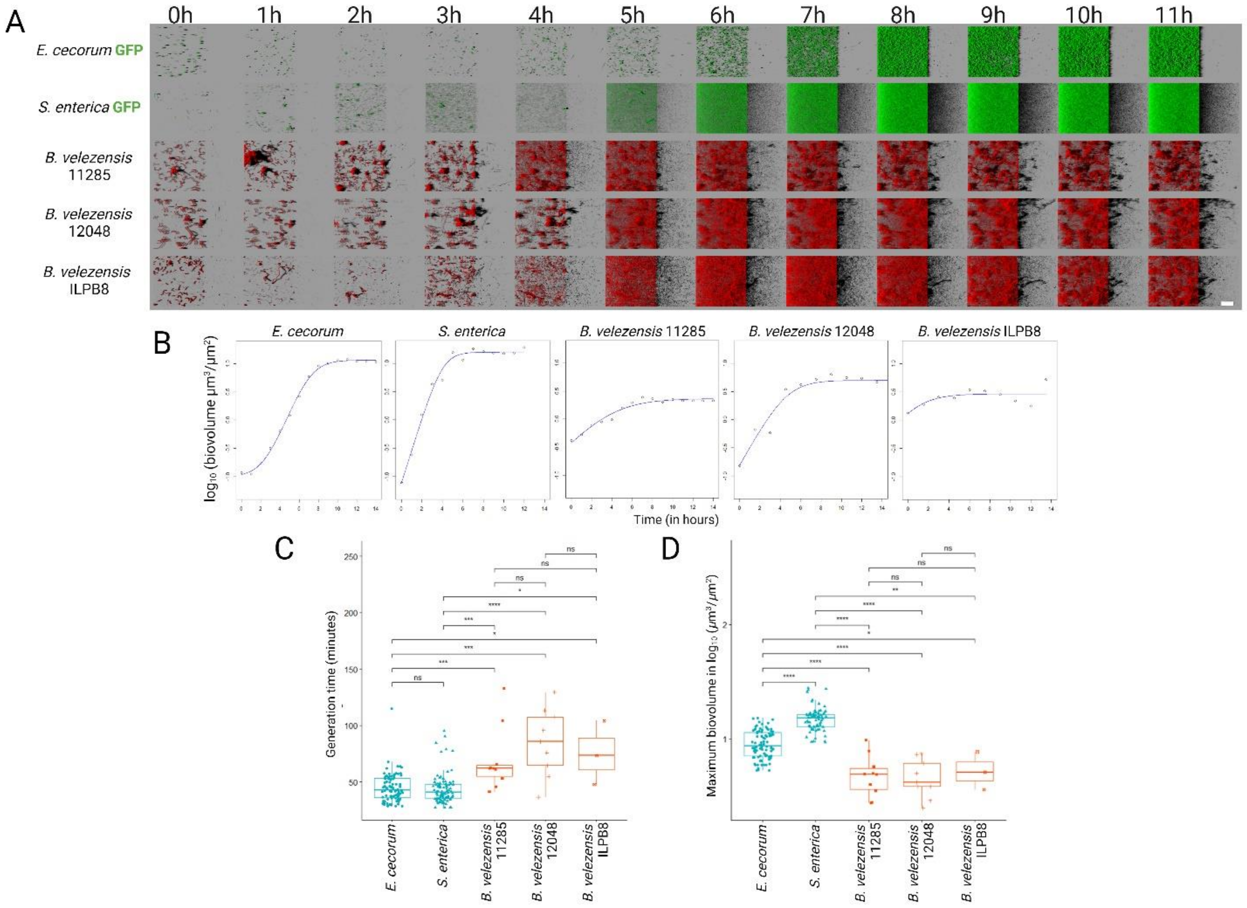


**Sup. 15 Biofilm formation dynamics through CLSM kinetic studies.** (A) Examples of CLSM visualisation of biofilm growth over time with a representative example for each control condition. The green colour corresponds to GFP labelling, while the red colour represents FM4-64. Scale bar = 40 µm. (B) Modelling of biofilm growth with one replicate per condition. The dots correspond to measured biovolumes, the line to the primary growth model fitted to observations. (C) Comparison of the generation times (D) and maximum biovolumes extracted using the mathematical model reached during monoculture experiments.

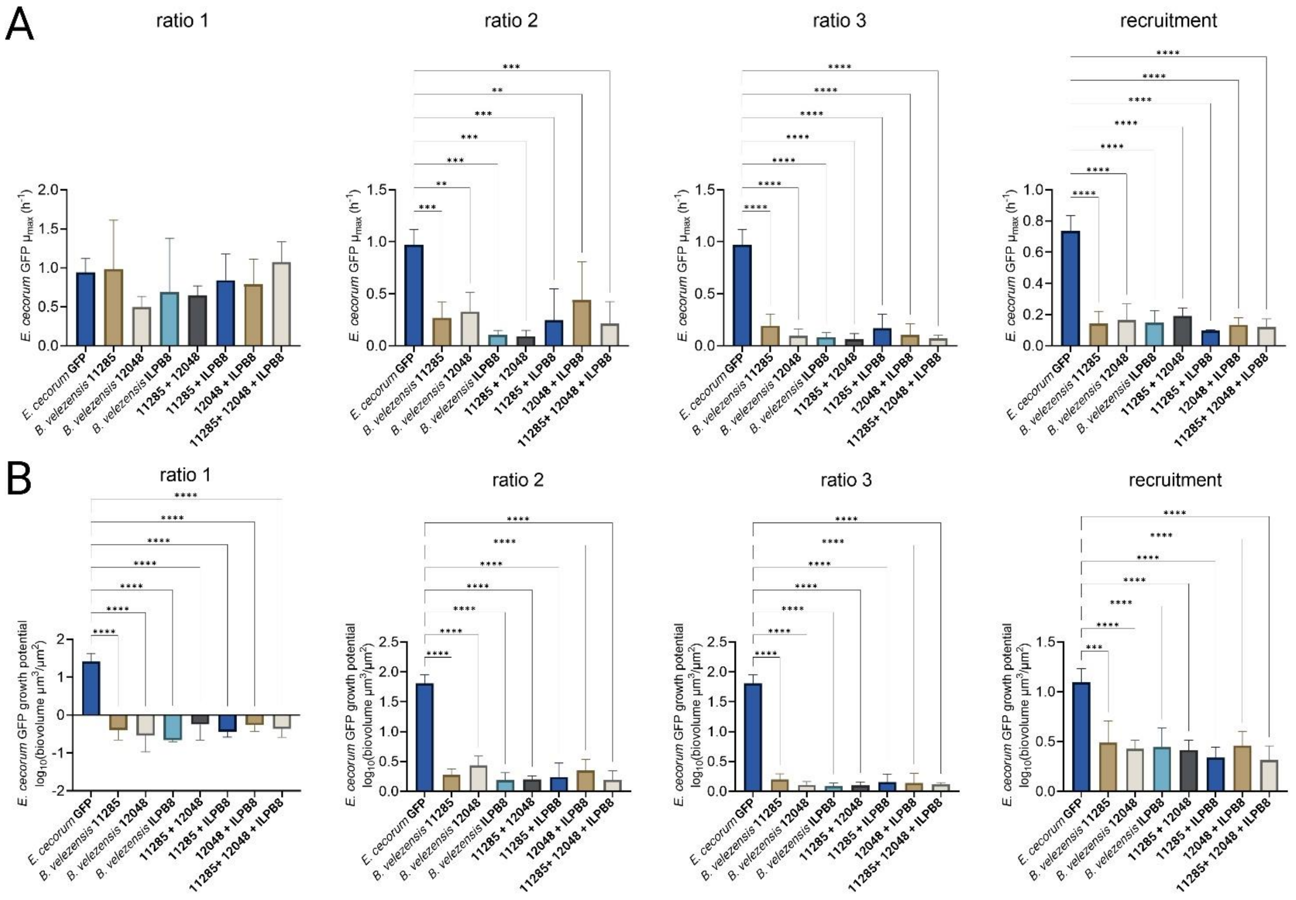


**Sup. 16 Growth rates and growth potentials of *E. cecorum* in the presence of *B. velezensis* alone or in consortia.** The initial biovolume ratios of *E. cecorum* GFP to *B. velezensis* were determined at the start of the experiment (ratio 1 = 1.4 (+/- 0.2), ratio 2 = 0.3 (+/- 0.06), ratio 3 = 0.03 (+/- 0.02), recruitment = 0.2 (+/- 0.04)). (A) Correspond to the growth rate and (B) to the growth potential.

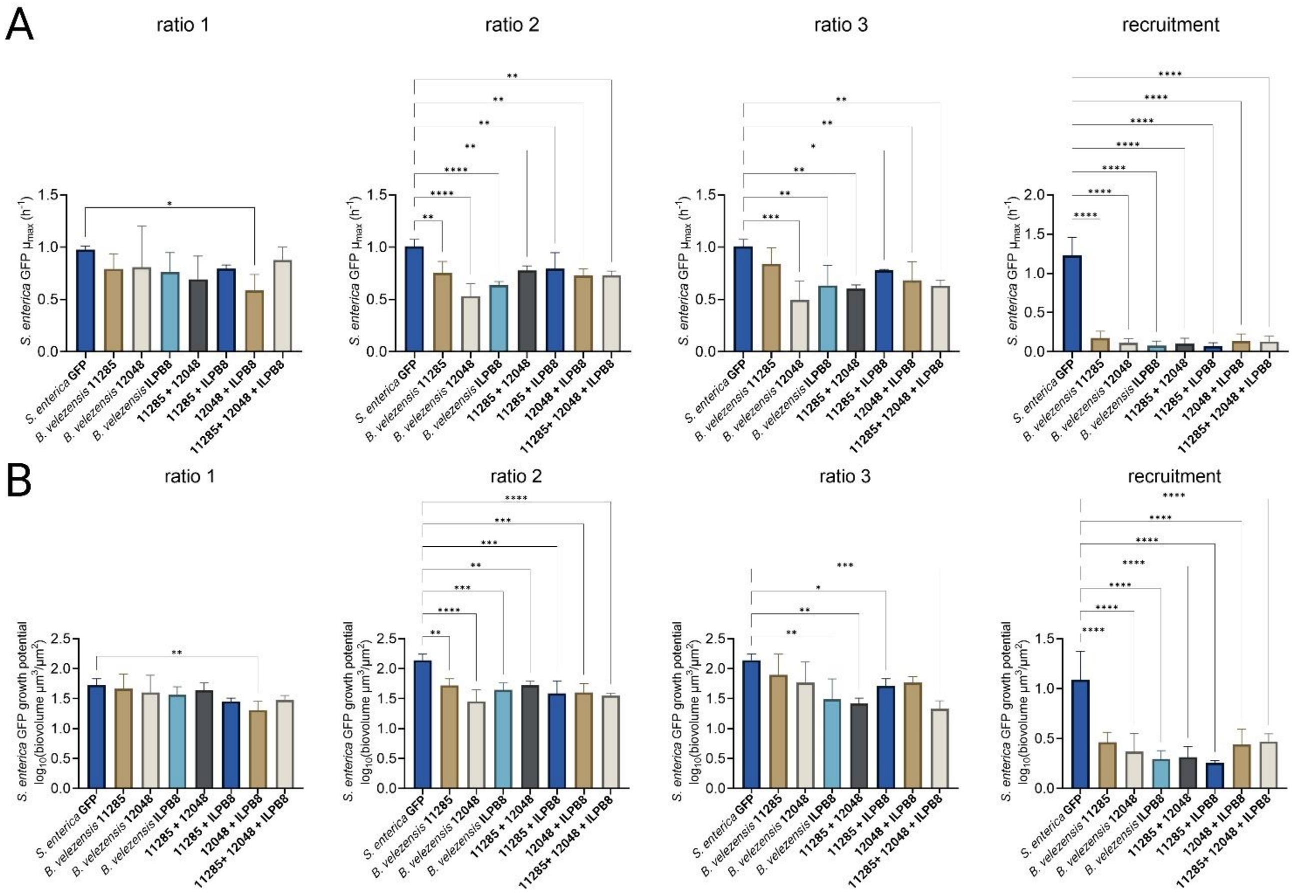


**Sup. 17 Growth rates and growth potentials of *S. enterica* in the presence of *B. velezensis* alone or in consortia.** The initial biovolume ratios of *S. enterica* GFP to *B. velezensis* were determined at the start of the experiment (ratio 1 = 3.2 (+/- 0.8), ratio 2 = 0.4 (+/- 0.1), ratio 3 = 0.1 (+/- 0.05), recruitment = 4.8 (+/- 0.8)). (A) Correspond to the growth rate and (B) to the growth potential.

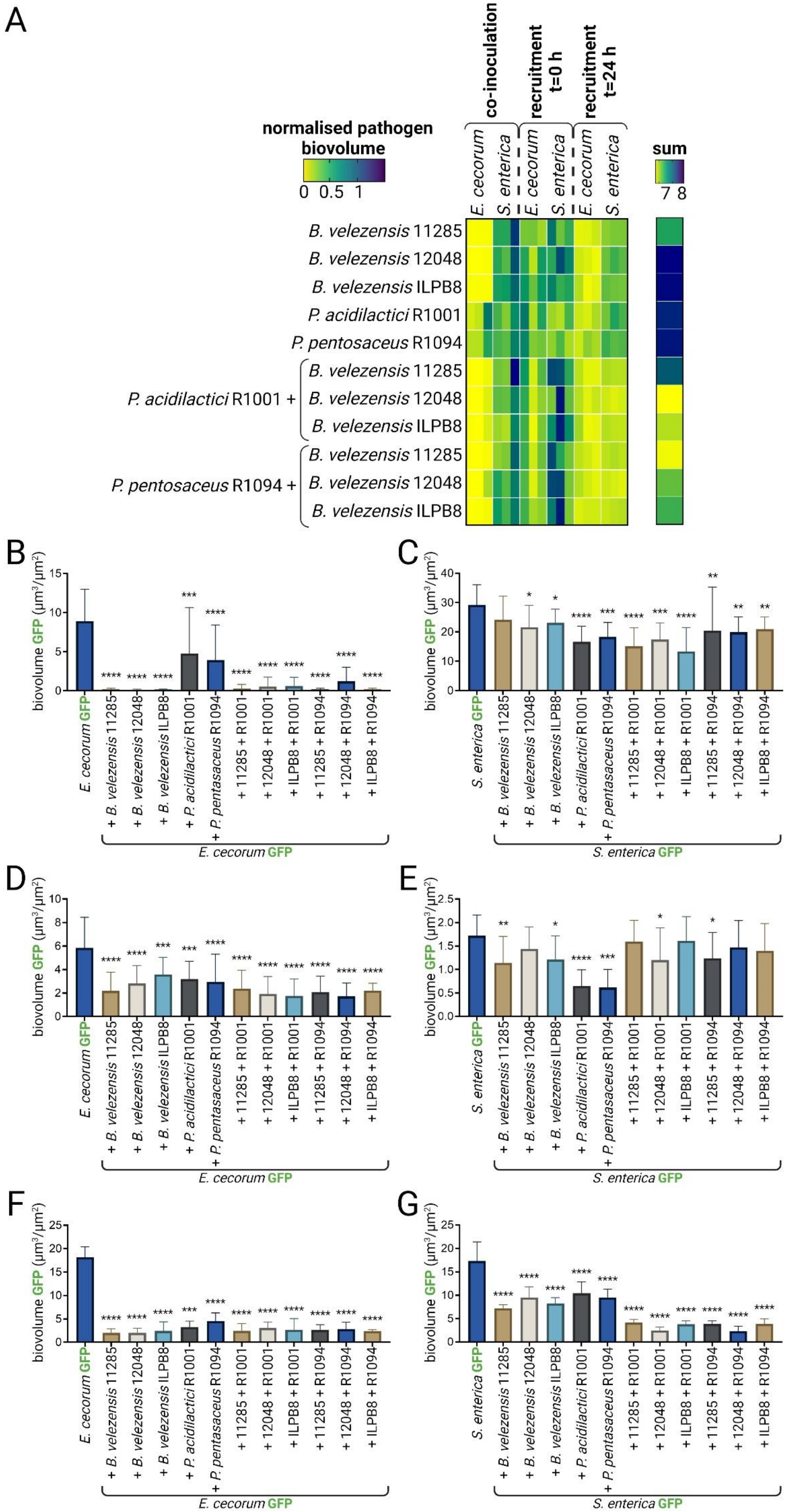
A
co-inoculation
recruitment t=0 h
recruitment t=24 h
normalised pathogen biovolume
0 0.5 1
sum
7 8
E. cecorum
S. enterica
E. cecorum
S. enterica
E. cecorum
S. enterica
B. velezensis 11285
B. velezensis 12048
B. velezensis ILPB8
P. acidilactici R1001
P. pentosaceus R1094
P. acidilactici R1001 +
B. velezensis 11285
B. velezensis 12048
B. velezensis ILPB8
P. pentosaceus R1094 +
B. velezensis 11285
B. velezensis 12048
B. velezensis ILPB8
B
C
D
E
F
G
biovolume GFP (µm³/µm²)
E. cecorum GFP
+ B. velezensis 11285
+ B. velezensis 12048
+ B. velezensis ILPB8
+ P. acidilactici R1001
+ P. pentasaceus R1094
+ 11285 + R1001
+ 12048 + R1001
+ ILPB8 + R1001
+ 11285 + R1094
+ 12048 + R1094
+ ILPB8 + R1094
S. enterica GFP

**Sup. 18 Exclusion of *E. cecorum* and *S. enterica* by *B. velezensis*, *Pediococcus* spp. or their combinations in the co-incubation models.** (A) A heatmap compiling all the results of GFP pathogen biovolumes in the presence of beneficial strains, normalised relative to the biovolumes of GFP pathogens alone. The detailed results are shown for (B) GFP-labelled *E. cecorum* in the co-inoculation model, (C) GFP-labelled *S. enterica* in the co-inoculation model, (D) GFP-labelled *E. cecorum* in the recruitment t=0h model, (E) GFP-labelled *S. enterica* in the recruitment t=0h model, (F) GFP-labelled *E. cecorum* in the recruitment t=24h model, (G) GFP-labelled *S. enterica* in the recruitment t=24h model. Error bars correspond to standard deviation. The biovolume values of the GFP pathogen in the presence of the candidate beneficial strains were compared with the biovolume of the pathogen alone.

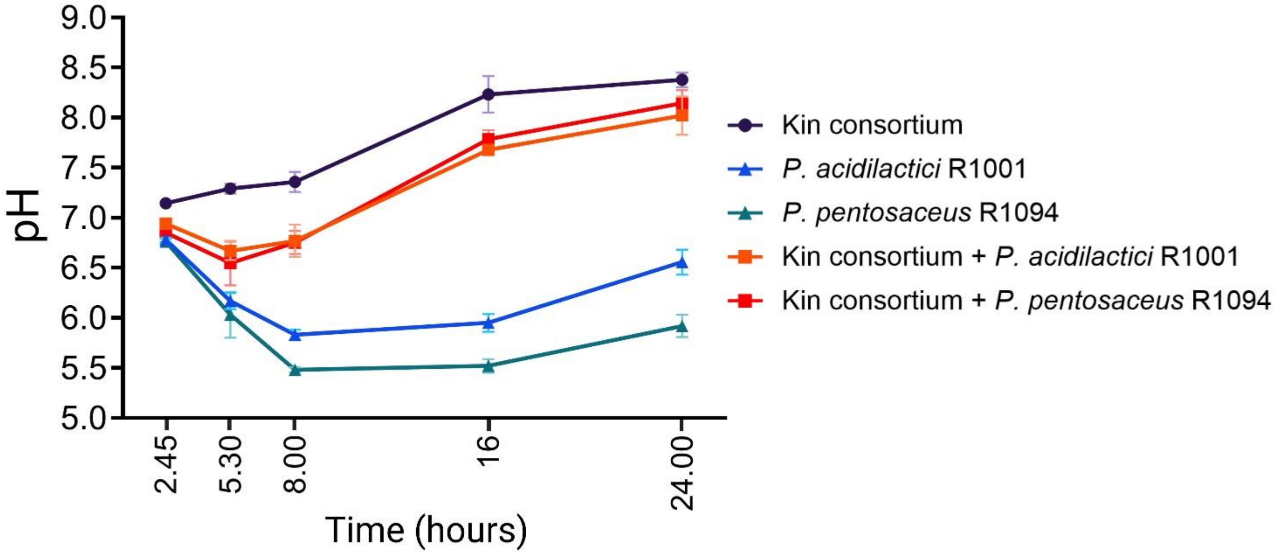


**Sup. 19 pH measurement of beneficial strains alone or in consortia, in the recruitment model.** The recruitment model was employed to monitor the pH of beneficial pathogen-free biofilm solutions over time. The volume equivalent to 18 wells of a 96-well plate (200 µL each, totalling 3.6 mL) was measured using a pH metre (Mettler Toledo, FiveEasy F20 model, France) for each biological replicate. Each measurement for a biological replicate corresponds to one sacrificed 96-well plate. Initially, bacteria adhered to the well bottoms following the co-inoculation protocol, and measurements were taken at 24 hours. Subsequently, the recruitment model was applied, where the medium was refreshed with fresh TS after co-inoculation, and measurements were conducted over time. Three biological replicates were performed for each measurement. The error bars represent standard errors.

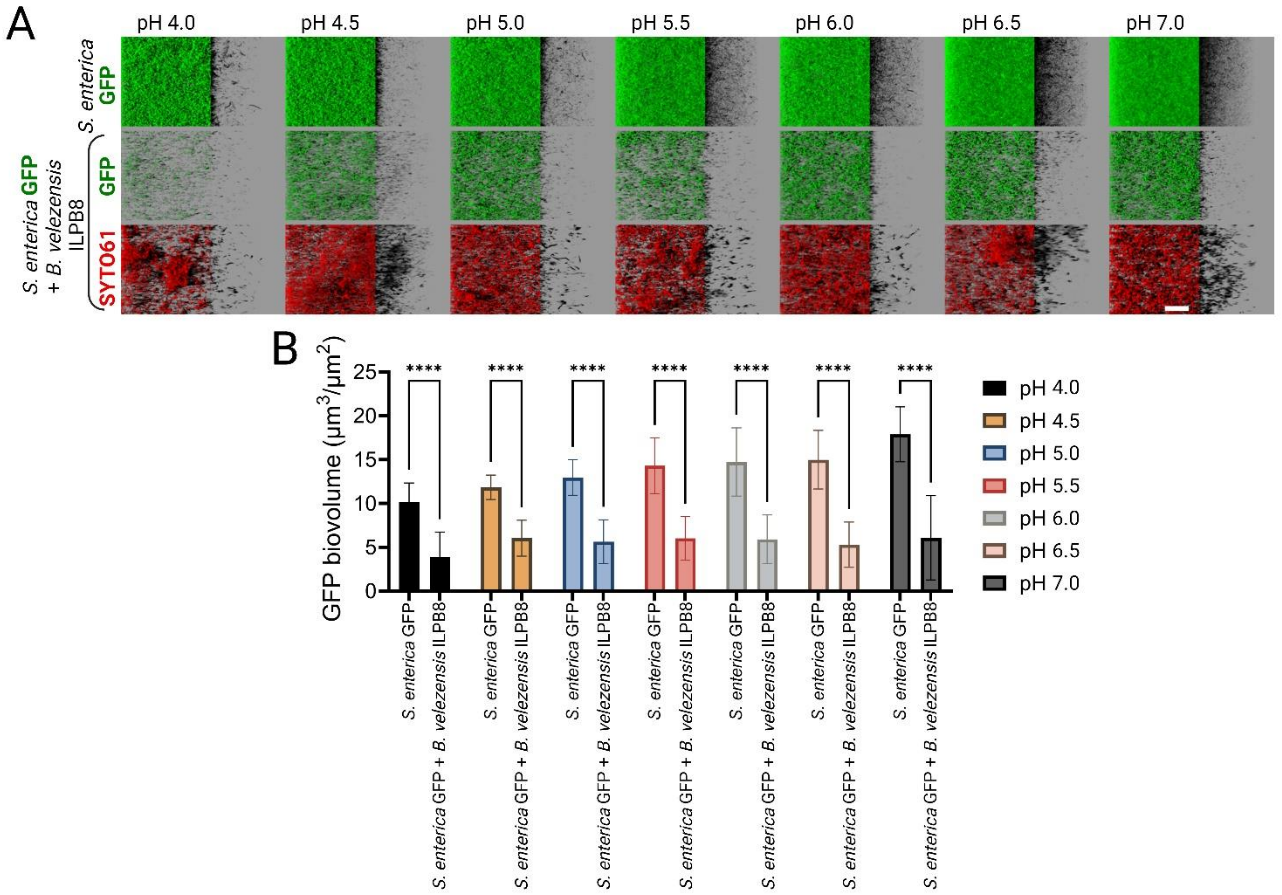


**Sup. 20 Biofilm growth of *S. enterica* with *B. velezensis* ILPB8 at different acidic pH values.** This experiment aims to investigate whether the pH reduction induced by *Pediococcus* spp. is involved in enhancing the exclusion of consortia composed of *Pediococcus* spp. and *B. velezensis* compared to *B. velezensis* alone, as observed in the recruitment model. *S. enterica* GFP is cultured either alone or in the presence of *B. velezensis* ILPB8 in the recruitment model at different acidic pH levels. HCl solution was incorporated to TSB medium to reach the desired value and then filtered with a 0.2 µm filter. *B. velezensis* biofilm is incubated for 24 hours, followed by the addition of *S. enterica* for a 1.5 h adhesion step before refreshing the media. (A) SYTO61 is added after 24 hours of incubation for observation using CLSM. (B) Through image analysis using the BiofilmQ software, the GFP biovolume of *S. enterica* alone is extracted and compared to the value obtained when *S. enterica* develops within a pre-established biofilm of *B. velezensis* ILPB8. The results demonstrate that medium acidification proportionally reduces the GFP biovolume of *S. enterica*. However, there is no significant difference in the *S. enterica* GFP biovolume in the presence of *B. velezensis* ILPB8 across different pH levels. Each value represents the average of three biological replicates, with four z-stack images each. The error bars correspond to standard errors.

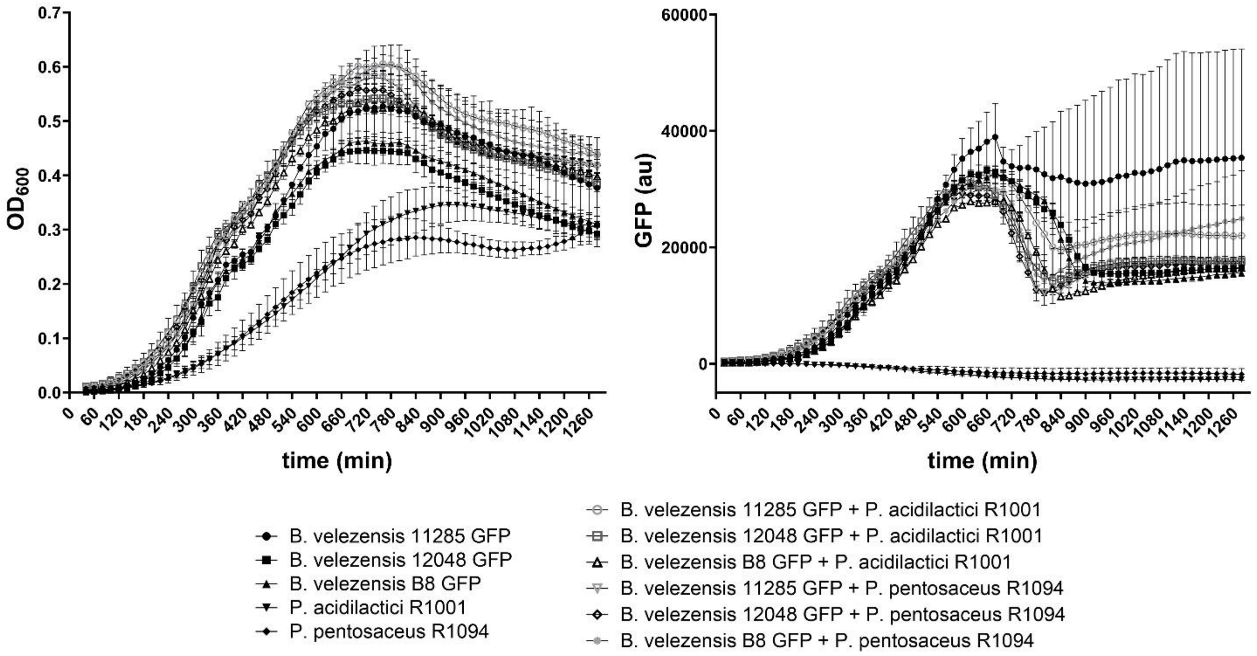


**Sup. 21 Incompatibility of mixtures of *B. velezensis* and *Pediococcus* spp. in a planktonic lifestyle.** The compatibility of *B. velezensis* expressing GFP and *Pediococcus* spp. Was investigated using the same protocol as the co-inoculation model. Following the adhesion step, ensuring the same initial biovolume for both bacterial partners, the cultures were resuspended. The cultures were continuously agitated to prevent bacterial biofilm formation. The OD600 was monitored over time using a Biotek plate reader (BioTek synergy h1, Agilent Technologies, USA) to follow the whole population. Fluorescence intensity between 500 and 550 was measured to assess the growth of *B. velezensis* GFP. Error bar represents SEM.

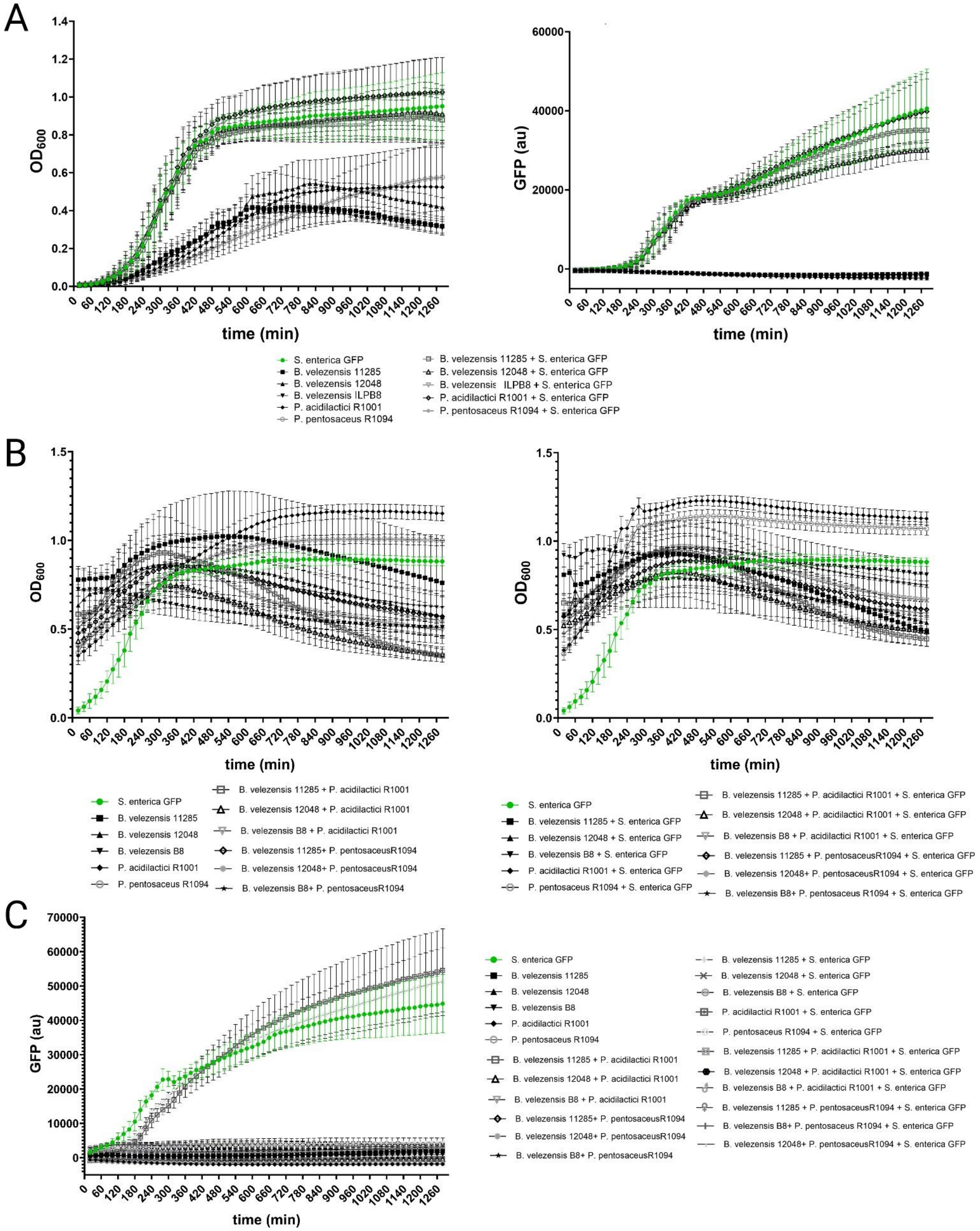
A
OD600
time (min)
GFP (au)
time (min)
S. enterica GFP
B. velezensis 11285
B. velezensis 12048
B. velezensis ILPB8
P. acidilactici R1001
P. pentosaceus R1094
B. velezensis 11285 + S. enterica GFP
B. velezensis 12048 + S. enterica GFP
B. velezensis ILPB8 + S. enterica GFP
P. acidilactici R1001 + S. enterica GFP
P. pentosaceus R1094 + S. enterica GFP
B
OD600
time (min)
OD600
time (min)
S. enterica GFP
B. velezensis 11285
B. velezensis 12048
B. velezensis B8
P. acidilactici R1001
P. pentosaceus R1094
B. velezensis 11285 + P. acidilactici R1001
B. velezensis 12048 + P. acidilactici R1001
B. velezensis B8 + P. acidilactici R1001
B. velezensis 11285+ P. pentosaceusR1094
B. velezensis 12048+ P. pentosaceusR1094
B. velezensis B8+ P. pentosaceusR1094
S. enterica GFP
B. velezensis 11285 + S. enterica GFP
B. velezensis 12048 + S. enterica GFP
B. velezensis B8 + S. enterica GFP
P. acidilactici R1001 + S. enterica GFP
P. pentosaceus R1094 + S. enterica GFP
B. velezensis 11285 + P. acidilactici R1001 + S. enterica GFP
B. velezensis 12048 + P. acidilactici R1001 + S. enterica GFP
B. velezensis B8 + P. acidilactici R1001 + S. enterica GFP
B. velezensis 11285 + P. pentosaceusR1094 + S. enterica GFP
B. velezensis 12048+ P. pentosaceusR1094 + S. enterica GFP
B. velezensis B8+ P. pentosaceus R1094 + S. enterica GFP
C
GFP (au)
time (min)
S. enterica GFP
B. velezensis 11285
B. velezensis 12048
B. velezensis B8
P. acidilactici R1001
P. pentosaceus R1094
B. velezensis 11285 + P. acidilactici R1001
B. velezensis 12048 + P. acidilactici R1001
B. velezensis B8 + P. acidilactici R1001
B. velezensis 11285+ P. pentosaceusR1094
B. velezensis 12048+ P. pentosaceusR1094
B. velezensis B8+ P. pentosaceusR1094
B. velezensis 11285 + S. enterica GFP
B. velezensis 12048 + S. enterica GFP
B. velezensis B8 + S. enterica GFP
P. acidilactici R1001 + S. enterica GFP
P. pentosaceus R1094 + S. enterica GFP
B. velezensis 11285 + P. acidilactici R1001 + S. enterica GFP
B. velezensis 12048 + P. acidilactici R1001 + S. enterica GFP
B. velezensis B8 + P. acidilactici R1001 + S. enterica GFP
B. velezensis 11285 + P. pentosaceusR1094 + S. enterica GFP
B. velezensis B8+ P. pentosaceus R1094 + S. enterica GFP
B. velezensis 12048+ P. pentosaceusR1094 + S. enterica GFP

**Sup. 22 Competition between *B. velezensis* or *Pediococcus* spp. and *S. enterica* in a planktonic lifestyle using different inoculation ratios.** Following the adhesion step to validate the desired biovolume for both partners at the start of the experiment, the bacteria were resuspended and the cultures continuously agitated to prevent biofilm formation. The OD600 was monitored over time using a BioTek plate reader (Biotek Synergy H1, Agilent Technologies, USA) to track the entire population. Fluorescence intensity between 500 and 550 was measured to assess the growth of S. enterica GFP. The error bars represent the standard error of the mean (SEM). (A) The protocol used for the co-inoculation model was employed to investigate the antagonistic effect of *B. velezensis* and *Pediococcus* spp. on S. enterica expressing GFP. The left graph represents the measured OD600, and the right graph shows GFP intensity. (B) OD600 growth of individual strains is shown on the left graph or in the presence of *S. enterica* GFP on the right graph. (C) GFP intensity in the presence of beneficial competitor strains. In this last graph, a growth of *S. enterica* GFP is only observed with *Pediococcus* spp. strains.

### Supplementary references


1. Guéneau V, Rodiles A, Frayssinet B *et al.* Positive biofilms to control surface-associated microbial communities in a broiler chicken production system - a field study. *Front Microbiol* 2022;**13**:981747.

2. Malone CL, Boles BR, Lauderdale KJ *et al.* Fluorescent reporters for Staphylococcus aureus. *J Microbiol Methods* 2009;**77**:251–60.

3. Olson RD, Assaf R, Brettin T *et al.* Introducing the Bacterial and Viral Bioinformatics Resource Center (BV-BRC): a resource combining PATRIC, IRD and ViPR. *Nucleic Acids Res* 2023;**51**:D678–89.

4. Prajsnar TK, Renshaw SA, Ogryzko NV *et al.* Zebrafish as a novel vertebrate model to dissect enterococcal pathogenesis. *Infect Immun* 2013;**81**:4271–9.
